# Designed miniproteins potently inhibit and protect against MERS-CoV

**DOI:** 10.1101/2024.11.03.621760

**Authors:** Robert J. Ragotte, M. Alejandra Tortorici, Nicholas J. Catanzaro, Amin Addetia, Brian Coventry, Heather M. Froggatt, Jimin Lee, Cameron Stewart, Jack T. Brown, Inna Goreshnik, Jeremiah N. Sims, Lukas F. Milles, Basile I.M. Wicky, Matthias Glögl, Stacey Gerben, Alex Kang, Asim K. Bera, William Sharkey, Alexandra Schäfer, Ralph S. Baric, David Baker, David Veesler

## Abstract

Middle-East respiratory syndrome coronavirus (MERS-CoV) is a zoonotic pathogen with 36% case-fatality rate in humans. No vaccines or specific therapeutics are currently approved to use in humans or the camel host reservoir. Here, we computationally designed monomeric and homo-oligomeric miniproteins binding with high affinity to the MERS-CoV spike (S) glycoprotein, the main target of neutralizing antibodies and vaccine development. We show that these miniproteins broadly neutralize a panel of MERS-CoV S variants, spanning the known antigenic diversity of this pathogen, by targeting a conserved site in the receptor-binding domain (RBD). The miniproteins directly compete with binding of the DPP4 receptor to MERS-CoV S, thereby blocking viral attachment to the host entry receptor and subsequent membrane fusion. Intranasal administration of a lead miniprotein provides prophylactic protection against stringent MERS-CoV challenge in mice motivating future clinical development as a next-generation countermeasure against this virus with pandemic potential.

## Introduction

Middle-East respiratory syndrome coronavirus (MERS-CoV) is a betacoronavirus causing severe and often deadly respiratory disease that was first identified in Saudi Arabia in 2012*(1, 2)*. MERS-CoV is a zoonotic virus, and most human cases result from direct viral transmission from the dromedary camel host reservoir or from human-to-human transmission in healthcare settings. Between April 2012 and August 2024, the European Centre for Disease Prevention and Control reported a total of 2,622 laboratory-confirmed MERS cases and 953 deaths globally, corresponding to a fatality rate of 36%*(3)*. MERS-CoV is endemic in camels in the Arabian Peninsula*(4)* and has recently been detected in humans in Africa*(5)*, underscoring possibly wider distribution and transmission than previously appreciated. To date, no vaccines or specific therapeutics are approved to prevent or treat MERS-CoV infections in humans or animals.

Entry of MERS-CoV into susceptible cells is mediated by the spike (S) glycoprotein which forms homotrimers protruding from the viral envelope*(6–8)*. S is the major antigen present at the viral surface and is the target of neutralizing antibodies during infection as well as the focus of vaccine design*(7, 9–11)*. MERS-CoV S comprises an S_1_ subunit, which mediates binding to sialosides via the N-terminal domain (NTD) and to the dipeptidyl-peptidase 4 entry receptor (DPP4) via the receptor-binding domain (RBD), as well as an S_2_ subunit which promotes viral and cellular membrane fusion to initiate infection*(7, 12–17)*. The differential distribution of these receptors in humans (lower airways) and camels (upper airways) possibly explains the severity but limited transmissibility of MERS-CoV in humans and the milder but more infectious profile in the reservoir host*(14, 18)*. The continued circulation of MERS-CoV and related merbecoviruses*(19–23)* and the risk of emergence of more transmissible strains motivate the development of potent and scalable therapeutics for pandemic preparedness. Computationally-designed miniproteins are emerging as competitive alternatives to monoclonal antibodies due to their high binding affinity, cost-effective manufacturing and the possibility to administer them directly in the upper respiratory tract (i.e. the initial site of infection for respiratory pathogens). Computational design of miniproteins targeting the SARS-CoV-2 RBD yielded potent and protective inhibitors of SARS-CoV-2 that retained activity against emerging SARS-CoV-2 variants throughout the COVID-19 pandemic*(24–27)*. Here, we set out to computationally design miniproteins targeting the MERS–CoV RBD and characterized the molecular basis of binding and inhibition, neutralization potency and breadth and *in vivo* prophylactic protection.

## Results

### Design of MERS-CoV RBD-directed miniproteins

Given that RBD-directed antibodies account for most plasma neutralizing activity in humans previously infected with SARS-CoV-2 or MERS-CoV*(9, 28–30)*, we decided to target this domain to design MERS-CoV miniprotein inhibitors. We used Rosetta-based computational methods to target hydrophobic residues within the MERS-CoV S receptor-binding motif (RBM)- hDPP4 interface through docking of a three-helix bundle library using RifDock*(25, 31)* **(Fig 1A-B)**. FastDesign was subsequently used for sequence-design of these scaffolds followed by filtering based on contact molecular surface and Rosetta ΔΔG. The best interface motifs (according to ΔΔG) were grafted onto new backbones using MotifGraft*(32)* and designs that bound to the biotinylated MERS-CoV S EMC/2012 RBD *via* yeast-surface display were subsequently optimized using site-saturation mutagenesis to identify affinity-enhancing mutations **(Fig S1A-B)**. The process yielded three constructs, designated cb3, cb4 and cb6, with respective binding affinities for the MERS-CoV S EMC/2012 RBD of 3.7 nM, 61 nM and 44 nM, as determined by surface-plasmon resonance **(Fig 1C)**.

**Fig 1.**
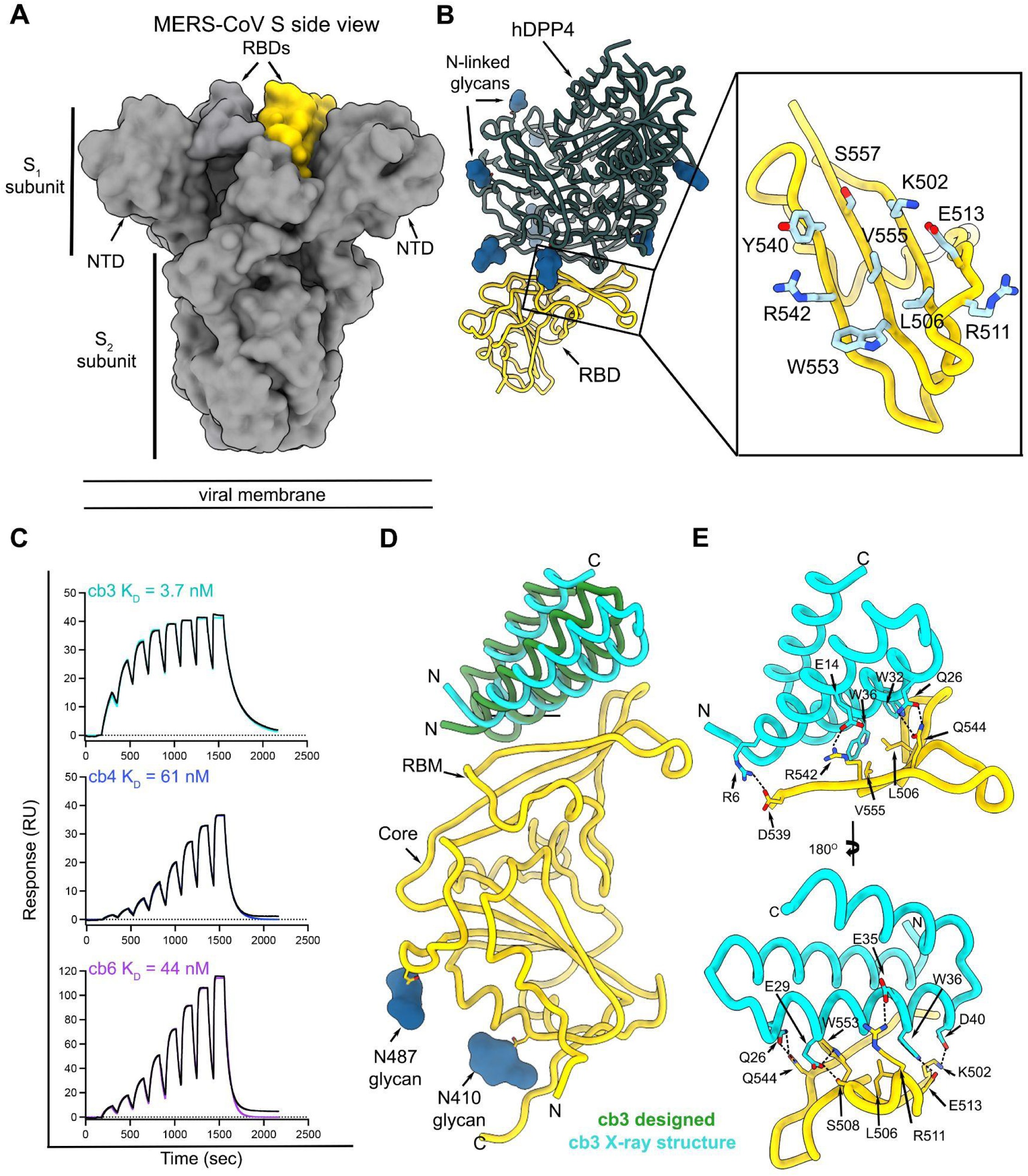
Computational design of MERS-CoV S RBD-targeting miniproteins. **A.** Surface representation of the prefusion MERS-CoV S trimer (grey, PDB 6Q04) highlighting the position of a single RBD in yellow. N-linked glycans are omitted for clarity. **B.** Ribbon representation of human DDP4 (dark green) in complex with the MERS-CoV S RBD (yellow) (PDB 4L72). Inset: zoomed-in view of the MERS-CoV S receptor-binding motif (RBM) highlighting the residues targeted for miniprotein design. **C.** Evaluation of binding of the cb3, cb4 and cb6 designed miniproteins to the immobilized MERS-CoV S EMC/2012 RBD using surface plasmon resonance (SPR). Eight miniprotein concentrations were used starting at 500 nM and following a two-fold dilution series. Data were analyzed using single cycle kinetics. RU: response units. Binding data and fits are shown as black and colored lines, respectively. **D.** Superimposition of the cb3-bound MERS-CoV S RBD (gold) from the computationally designed model (green) and the experimental crystal structure (cyan). Only one RBD is shown for clarity. **E.** Zoomed-in views of the crystal structure of the complex between cb3 (cyan) and the MERS-CoV S RBD (gold) highlighting selected interactions. Dashed lines indicate hydrogen bonds and salt bridges.

To characterize the molecular basis of MERS-CoV S recognition by these miniproteins, we determined a crystal structure of the MERS-CoV S RBD in complex with cb3 at 1.85 Å resolution using X-ray diffraction data **(Table S1)**. cb3 interacts with the MERS-CoV S RBM through polar interactions and shape complementarity burying an average surface of 750 Å^2^ at the interface between the two binding partners **(Fig 1D-E)**. cb3 utilizes its N-terminal two helices to interact with the MERS-CoV S RBD residues 502, 506-508, 511-513, 536-542, 544, 552-553, 555 without contacts with N-linked glycans. cb3 residues R6, E14, E35 and D40 are salt bridged to MERS-CoV S RBD residues D539, R542, R511 and K502, respectively, and a constellation of hydrogen bonds and van der Waals interactions strengthen the binding interface, explaining the single digit nanomolar affinity of this miniprotein **(Fig 1D-E)**. Superimposition of the MERS-CoV S RBD from the crystal structure determined here with that of the computationally designed complex reveals that the cb3 miniprotein aligns with an RMSD of 4.8 Å for 57 aligned Cα positions **(Fig 1F)**.

### Monovalent and multivalent miniproteins broadly neutralize MERS-CoV variants

To evaluate the neutralization potency and breadth of the monomeric miniproteins, we quantified their dose-dependent inhibitory activity using vesicular stomatitis virus (VSV) particles pseudotyped with a panel of MERS-CoV S variants. Our panel included the S glycoprotein of the following MERS-CoV isolates: EMC/2012 (NC_019843.3EMC/2012), United Kingdom/2012 (NC_038294.1), Jordan/2012 (AHY21469.1), Seoul/2015 (KT374056.1), and Kenya/2019 (OK094446.1) **(Table S2)**. All of these variants were identified in humans except for MERS-CoV Kenya/2019 which was detected in a camel*(33)*. Relative to EMC/2012 S, these variants comprise 2-8 amino acid mutations with United Kingdom/2012 S and Seoul/2015 S harboring residue changes known to reduce the neutralizing activity of monoclonal antibodies and polyclonal sera*(34–36)*. Out of the three monovalent miniproteins designed, only cb3 had detectable neutralizing activity against MERS-CoV EMC/2012 S VSV, likely due to differences in overall binding affinity which is an order of magnitude greater for cb3 relative to cb4 and cb6 **(Fig 2A-B, S2, Table S3)**. cb3 broadly inhibited MERS-CoV pseudoviruses harboring all S variants in our panel with 4.5-fold reduced potency for the United Kingdom/2012 isolate, compared to EMC/2012, most likely due to the L506F substitution within the cb3 binding site **(Fig 2A-B and S2)**.

**Fig 2.**
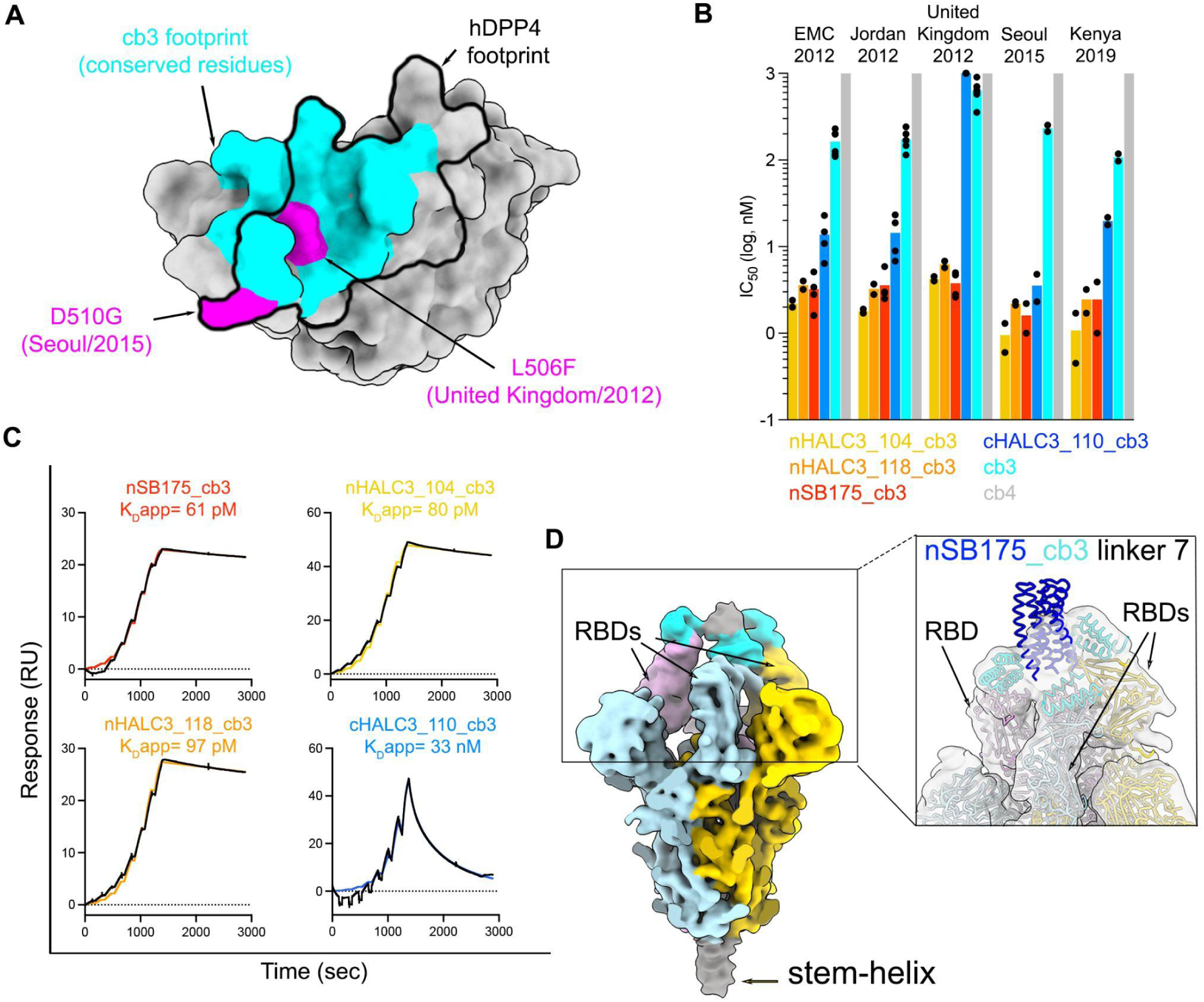
Miniprotein homotrimerization enhances neutralization potency. **A.** Surface representation of the MERS-CoV EMC/2012 RBD highlighting in cyan the conserved residues in the cb3-binding site among MERS-CoV EMC/2012, Jordan/2012, United Kingdom/2012, Seoul/2015 and Kenya/2019. The L506F and D510G residue substitutions present in the United Kingdom/2012 and Seoul/2015 isolates, respectively, are shown in pink whereas the hDPP4 footprint is shown as a black outline. **B.** Neutralizing activity of monomeric and trimeric miniproteins against a panel of MERS-CoV S variant VSV pseudoviruses. All miniproteins in this panel harbor a GSG linker between the oligomerization domain and the miniproteins. cb4, for which we did not detect neutralization, was plotted at the limit of detection (LOD) of 10^3^ nM. Each dot represents an IC_50_ expressed in nM obtained from a biological experiment. Bars represent the mean of the IC_50_s. Each biological experiment was performed with technical duplicates and curves are shown in Fig. S2, S3 and S4. The IC_50_s plotted here are listed in Fig S1, Tables S6 and S7. **C**. SPR analysis of binding of trimeric cb3 constructs to the prefusion biotinylated MERS-CoV EMC/2012 S ectodomain trimer immobilized on chips. Single cycle kinetic analysis of a two-fold dilution series starting at 25 nM is shown except for cHALC3_110_cb3 which had an upper concentration of 500 nM. Binding data and fits are shown as black and colored lines, respectively. **D.** CryoEM reconstruction of the nSB175_cb3 linker 7 bound to prefusion MERS-CoV- S low-pass filtered at 8 Å resolution and colored by protomer. The density corresponding to three bound cb3s is colored in cyan and the remaining density corresponding to the trimerization domain is colored gray. Inset: zoomed-in view of a model depicting nSB175_cb3 linker 7 (cb3-GGGSGGGS-SB175) fitted into the map shown in panel D (linkers were not modeled).

Based on our previous success in designing a homotrimeric, pan-variant SARS-CoV-2 inhibitor, designated TRI2-2*(24–27)*, we set out to engineer homotrimeric versions of cb3 to enable simultaneous engagement of all three RBDs within a MERS-CoV S trimer and enhance potency and breadth further through avidity. We developed multivalent miniproteins using a set of validated trimerization motifs*(24, 37)* fused to the cb3 N or C terminus *via* a GSG linker **(Fig S1)**. SPR analysis of binding of the resulting homotrimeric cb3 designs to immobilized prefusion MERS-CoV S showed that multimerized constructs with an N-terminal miniprotein and C-terminal oligomerization domain (denoted with prefix ‘n’ e.g. nHALC3_104_cb3) interacted with higher apparent affinities and had markedly reduced off-rates relative to monomeric cb3, consistent with avid binding **(Fig 2C and Table S3)**. Evaluation of neutralizing activity against the MERS-CoV EMC/2012, Jordan/2012 and United Kingdom/2012 pseudoviruses revealed that the oligomeric cb3 miniproteins were endowed with at least an order of magnitude greater potency, compared to monomeric cb3, with the exception of cHALC3_919_cb3, cHALC3_110_cb3, cHALC3_118_cb3, cHALC3_114_cb3 and cSB175_cb3 fusions which failed to neutralize MERS-CoV United Kingdom/2012 S VSV pseudotypes **(Fig 2B, Fig S3 and Table S6)**. We therefore selected the nHALC3_104_cb3, nHALC3_118_cb3 and nSB175_cb3 (cb3-GSG-SB175) homotrimeric miniproteins for further characterization of neutralization breadth and observed that all three of them inhibited the MERS-CoV Seoul/2015 and Kenya/2019 S VSV pseudoviruses, markedly outperforming monomeric cb3-mediated neutralization **(Fig 2B, Fig S4 and Table S7)**.

Storage and shipping conditions are key factors affecting the development of therapeutic molecules as they can impact the cost of manufacturing along with the practicality of distribution and access. Lyophilization has emerged as a method of storage and administration of protein biologics to increase shelf life and optimize *in vivo* release*(38, 39)*. Given that the SARS-CoV-2 S-directed TRI2-2 miniprotein is endowed with high biophysical stability, we set out to evaluate retention of activity of the nSB175_cb3 homotrimeric MERS-CoV miniprotein after lyophilization and reconstitution. We found that *in vitro* neutralization potency and breadth against our panel of MERS-CoV S variants were unaffected by freeze/thawing or lyophilization/reconstitution of nSB175_cb3, underscoring the optimal biophysical properties of this miniprotein **(Fig S5)**.

CryoEM characterization of nSB175_cb3 bound to prefusion MERS-CoV EMC/2012 S revealed that the observed enhancement of avidity and neutralizing activity afforded by homotrimerization results from engagement of two or three RBDs within each trimer **(Fig S6, Fig S7A-B and Table S8)**. We subsequently produced nSB175_cb3 homotrimeric constructs harboring systematic variations of linker length and evaluated their neutralizing activity against our panel of MERS-CoV S variants **(Fig S7C and Table S5)**. Given that the two constructs harboring the longest linkers, designated nSB175_cb3 linker 7 (cb3_GGGSGGGS_SB75) and nSB175_cb3 linker 8 (cb3_GGGSGGGSGGGS_SB75), exhibited slight improvements in neutralization potency **(Fig S6C),** we determined an asymmetric cryo-EM reconstruction of the former miniprotein construct bound to MERS-CoV S at 2.6 Å resolution, which also indicated variable minibinder stoichiometry among S trimers (**Fig 2D, Fig S8 and Table S8**). The quality of the cryo-EM density corresponding to the SB175 trimerization domain was reduced due to conformational heterogeneity caused by the flexible nature of the linker, however, subsequent local refinement of the region corresponding to two cb3 modules bound to two MERS-CoV S RBD yielded a reconstruction at 3.2 Å resolution that was virtually indistinguishable from the crystal structure of the monomeric cb3-bound MERS-CoV S RBD. These results indicate that oligomerization enabled retention of the binding mode designed (**Fig S8D**).

Collectively, these data show that the multimeric miniproteins potently inhibit MERS-CoV variants known to evade monoclonal antibody-mediated neutralization and are hyperstable, which are two highly desirable properties for the development of countermeasures against coronaviruses.

### The nSB175_cb3 (cb3-GSG-SB175) miniprotein interferes with MERS-CoV S binding to the host receptor

Our structural data suggest that cb3 binding to the MERS-CoV RBD would be incompatible with engagement of the hDPP4 receptor due to steric overlap **(Fig 3A)**. To test this hypothesis we assessed whether nSB175_cb3 and hDPP4 can simultaneously bind to the MERS-CoV RBD using biolayer interferometry. We found that nSB175_cb3 binding to the immobilized MERS-CoV RBD blocked subsequent binding of hDPP4, confirming the expected competition **(Fig 3B)**. We next investigated whether the nSB175_cb3 homotrimer impacted cell-cell fusion mediated by the MERS-CoV EMC/2012, United Kingdom/2012, Jordan/2012, Seoul/2015, and Kenya/2019 S variants. We recorded live cell imaging using a split green fluorescent protein (GFP) system using VeroE6 target cells stably expressing human TMPRSS2 and GFP_1-10_ and BHK-21 effector cells stably expressing GFP_11_ and transiently transfected with a MERS-CoV S variant*(40)*. Analysis of S cell-surface expression levels by flow cytometry showed consistently lower expression of MERS-CoV S EMC/2012, Seoul/2015 and Kenya/2019 relative to Jordan/2012 and United Kingdom/2012 (**Fig S9A-B**), which was compensated for by adjusting the amount of cells to yield similar fusion readings. We monitored cell-cell fusion over a period of 18 h in the presence of different concentrations of nSB175_cb3 and observed dose-dependent inhibition of membrane fusion which was completely abrogated at a concentration of 660 nM **(Fig 3 C-D and Fig S9C)**. The SARS-CoV-2 S-directed TRI2-2 miniprotein, which uses the same trimerization domain*(24)*, had no effect on fusion, confirming the specificity of inhibition observed **(Fig 3C)**. These data indicate that the broad neutralizing activity of nSB175_cb3 against MERS-CoV S variants results from direct competition with viral attachment to the host cell receptor DDP4, which in turn blocks membrane fusion.

**Fig 3.**
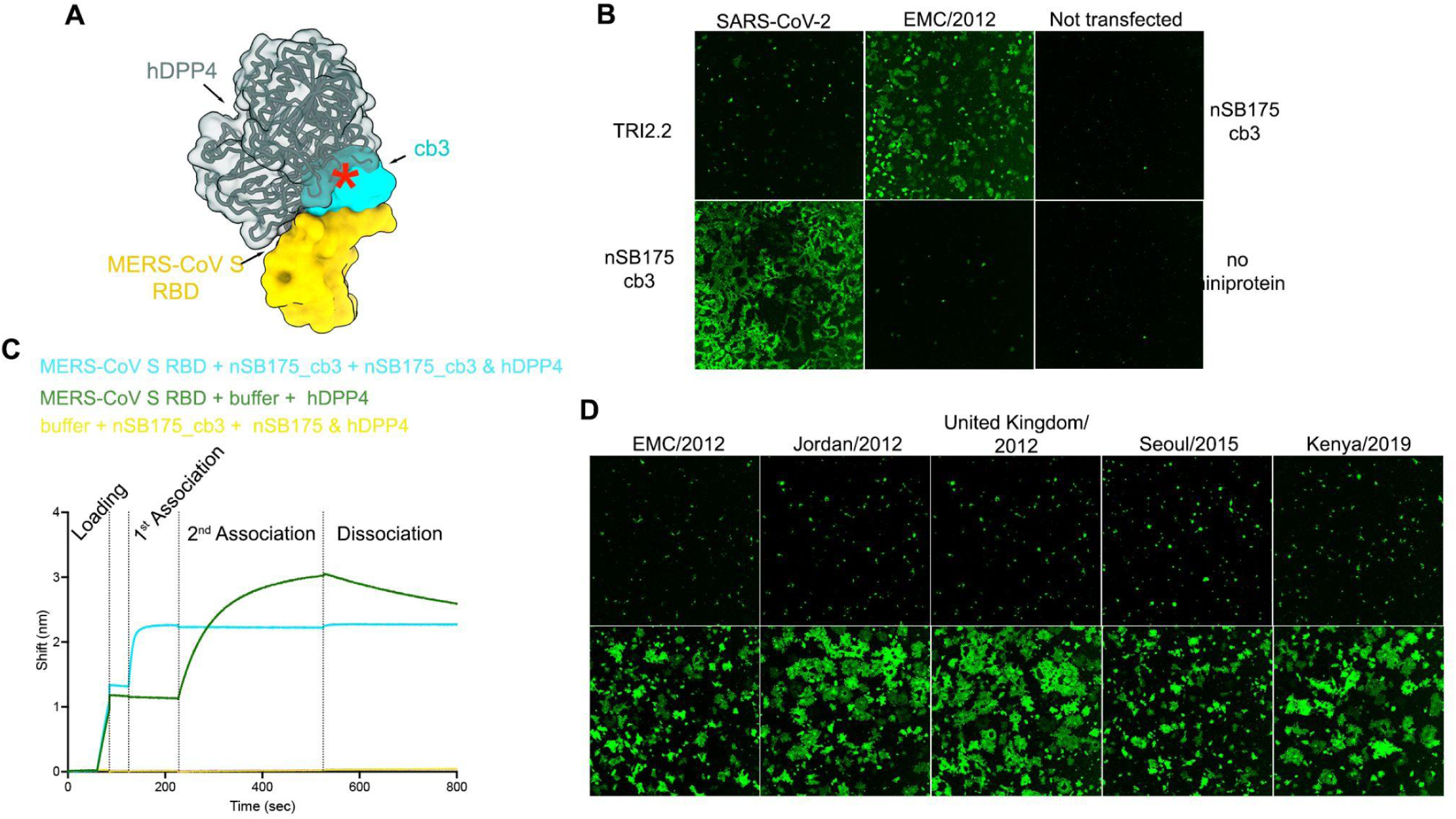
The homotrimeric nSB175_cb3 (cb3-GSG-SB175) miniprotein inhibits MERS-CoV attachment to the hDPP4 receptor. **A.** Composite model showing that the miniprotein cb3 (cyan surface) and human DPP4 (hDPP4, dark green) bind to partially overlapping binding sites on the MERS-CoV RBD (rendered as a yellow surface). The red asterisk indicates steric clashes. **B.** BLI analysis of hDPP4 binding to MERS-CoV S RBD. Streptavidin biosensor associated with biotinylated MERS-CoV S RBD were dipped in a solution containing either 0.5 µM of nSB175_cb3 or buffer (1^st^ association) and subsequently in a solution containing 0.5 µM of hDPP4 and 0.5 µM of nSB175_cb3 (cyan) or 0.5 µM of hDPP4 (green) (2^nd^ association). Streptavidin biosensors with no MERS-CoV S RBD associated were use as a negative control (yellow) **C**. nSB175_cb3-mediated inhibition (at a concentration of 660 nM) of cell-cell fusion between BHK-21 cells transiently transfected with MERS-CoV (EMC/2012) S or SARS-CoV-2 (Wuhan-Hu-1) S and VeroE6-TMPRSS2 followed by reconstitution of a split GFP. Untransfected cells were used as negative control. SARS-CoV-2 TRI2.2 is a SARS-CoV-2 S-specific miniprotein. **D**. nSB175_cb3-mediated inhibition (at a concentration of 660 nM) of cell-cell fusion mediated by the indicated MERS-CoV S variants in conditions identical to (B). Images in C and D correspond to the end point of an 18-hour time course experiment.

### nSB175_cb3 (cb3-GSG-SB175) protects mice against MERS-CoV challenge

To evaluate the *in vivo* protective efficacy of one of these homotrimeric miniproteins, we inoculated susceptible C57BL/6J 288/230 mice*(41)* with 10^5^ p.f.u. MERS-CoV-m34c5 intranasally and followed weight loss as a proxy for disease for 5 days. nSB175_cb3 was administered intranasally at 6.25 mg/kg one day before challenge and control groups were treated identically except that they received an influenza virus hemagglutinin-directed miniprotein, buffer alone (TBS), or were untreated **(Figure 4A, S10)**. Prophylactic administration of nSB175_cb3 completely protected challenged mice against weight loss throughout the duration of the experiments and abrogated viral replication in the lungs. In contrast, control groups experienced 10-20% weight loss and had ∼3 orders of magnitude greater lung viral titers, relative to animals receiving nSB175_cb3 **(Figure 4 B-C)**. Gross pathological assessment of lung tissues at 5 days post viral challenge revealed that nSB175_cb3 prophylaxis entirely protected from visible signs of lung hemorrhage, as opposed to control groups that uniformly exhibited lung discoloration induced by MERS-CoV **(Figure 4D)**. Mice treated with nSB175_cb3 pre- and post-MERS-CoV challenge were protected from weight loss, lacked gross pathological lung abnormalities and had significantly reduced viral titres in the lung, mirroring the results of prophylactic administration alone (**Figure S10**). These data show that a single dose of nSB175_cb3 administered one day prior to viral exposure provides prophylactic protection against a stringent MERS-CoV challenge and reduces lung viral burden.

**Fig 4.**
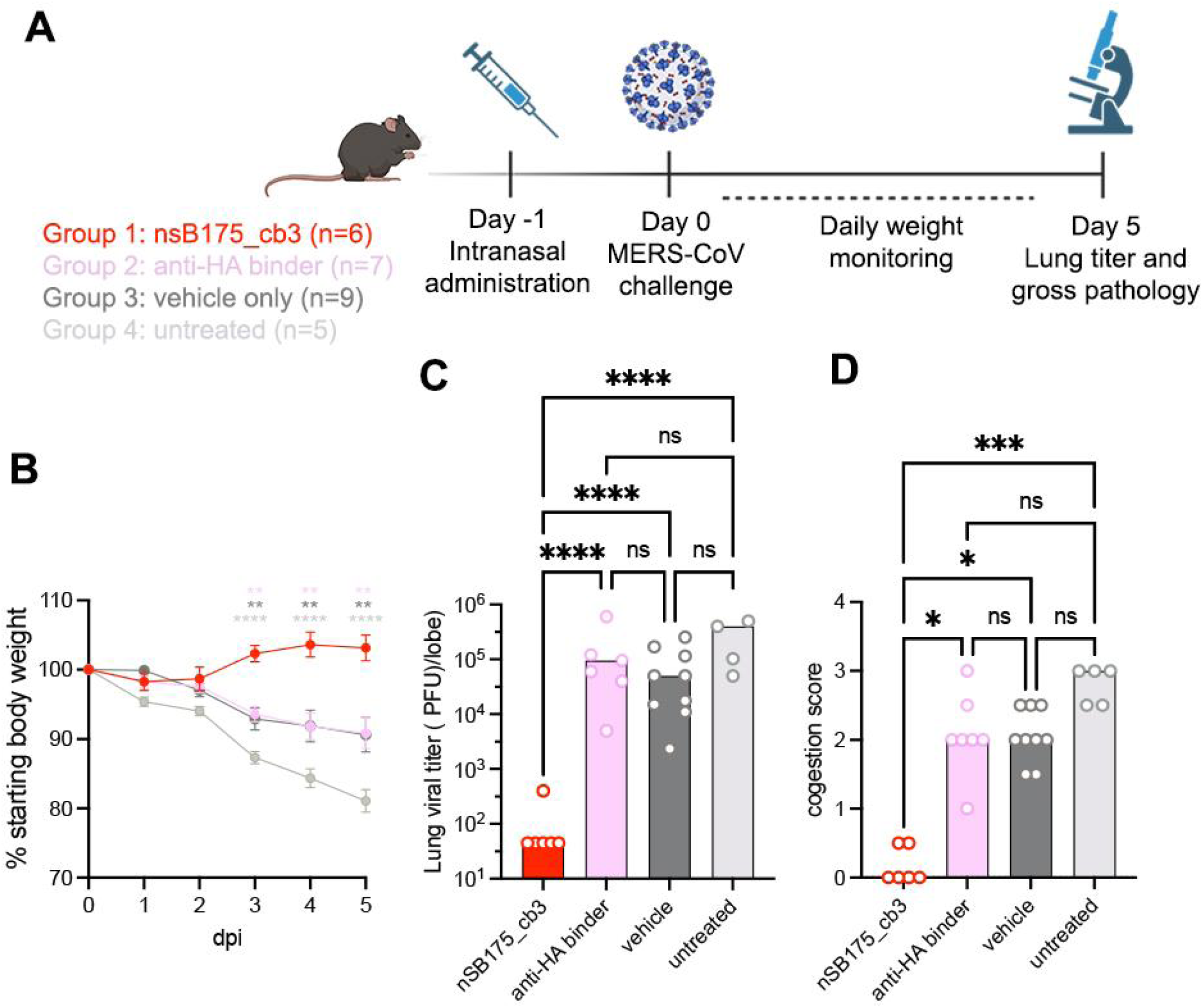
Intranasal administration of nSB175_cb3 (cb3-GSG-SB175) provides prophylactic protection against MERS-CoV challenge. **A.** nSB175_cb3, monomeric influenza hemagglutinin miniprotein (anti-HA) or buffer alone (TBS) were administered intranasally to C57BL/6 J288/330 mice one day prior to challenge with MERS-CoV-m34c5. Animals were monitored for 5 days for body weight and analyzed at day 5 for congestion score and replicating viral titer in lungs. **B**. Weight loss at different days post infection (dpi). **C**. Lung viral titers at day 5 post infection.￼ **D**. Congestion score 5 days post infection. Group comparisons for body weight and viral titers were assessed with the two-way ANOVA: Tukey’s test. For the congestion score, comparisons among groups were assessed with the Kruskal-Wallis test; ns, not significant; *P < 0.05, ***P < 0.001, ****P < 0.0001.

## Discussion

A comprehensive pandemic preparedness strategy should include the development of medical interventions that can rapidly curtail transmission and lethality of priority pathogens during outbreaks, as opposed to relying on new medicines developed only once the pandemic has started. Emergency drug-use authorizations took months to years to be granted during the COVID-19 pandemic for prophylactic and therapeutic monoclonal antibodies developed against SARS-CoV-2*(11, 42)*. An ideal MERS-CoV antiviral would inhibit a wide diversity of MERS-CoV variants, protect against disease, limit transmission, enable cost-effective and scalable manufacturing and distribution, while having a minimally invasive route of administration. Intranasally administered countermeasures are especially attractive as they are delivered directly at the site of initial viral infection and they can act immediately, without delays to mount an immune response as is the case for vaccines.

The nSB175_cb3 miniprotein MERS-CoV inhibitor developed in this work is an excellent candidate for such pre-pandemic MERS antiviral development. nSB175_cb3 blocks interactions of MERS-CoV S with the DPP4 entry receptor, and provides prophylactic protection against disease by reducing viral burden in the lungs by three orders of magnitude. The broadly neutralizing activity of nSB175_cb3 against a panel of MERS-CoV S variants demonstrates some degree of resilience to viral evolution and antigenic changes, as that observed for SARS-CoV-2 variants throughout the COVID-19 pandemic*(40, 43–47)*. Given the threat that MERS-CoV poses to global biosecurity, the continued clinical development of this molecule should assess its ability to accelerate recovery when administered solely after exposure as well as its ability to limit transmission. The development of viral inhibitors, assessment of their safety, manufacturing and stockpiling prior to the emergence of a pandemic variant of a given virus is a compelling approach to ensure medical countermeasures are available for emerging and re-emerging threats.

## Acknowledgements

RJR is a Washington Research Foundation Postdoctoral Fellow. This study was supported by the National Institute of Allergy and Infectious Diseases (NIHAID) (AI171292 to RSB, R01AI160052 to D.B. and D.V., P01AI167966, DP1AI158186 and 75N93022C00036 to D.V.), an Investigators in the Pathogenesis of Infectious Disease Awards from the Burroughs Wellcome Fund (D.V.), the University of Washington Arnold and Mabel Beckman cryoEM center and the National Institute of Health grant S10OD032290 (to D.V.). D.B. and D.V. are Investigators of the Howard Hughes Medical Institute. D.V. is the Hans Neurath Endowed Chair in Biochemistry at the University of Washington. The X-ray crystallography data of MERS-CoV RBD and cb3 complex was remotely collected at the National Synchrotron Light Source II AMX beamline. The HALC3_919 crystallographic data was collected at the Northeastern Collaborative Access Team beamlines at Advanced Photon Source, which are funded by the National Institute of General Medical Sciences from the National Institutes of Health (P30 GM124165). This research used resources of the Advanced Photon Source, a U.S. Department of Energy (DOE) Office of Science User Facility operated for the DOE Office of Science by Argonne National Laboratory under Contract No. DE-AC02-06CH11357. Quantification of the different MERS-CoV S surface expressed on BHK-21-GFP_1-10_ cells was performed at the Flow Cytometry Core from the Department of Laboratory Medicine and Pathology (DLMP) at the University of Washington.

## Author Contributions

MAT, RJR, BC, DB, and DV conceived the study and designed the experiments; RJR and BC designed the monomeric and trimeric miniproteins with support from BW and LM. IG did the yeast surface display experiments with support from JS. MAT carried out X-ray crystallography and cryoEM work of MERS-CoV RBD and S complexes with help from DV. MAT and RJR performed binding studies with support from MG. MAT, CS, JTB and WS recombinantly expressed and purified glycoproteins. RJR and SG recombinantly expressed and purified designed binders. RJR carried out X-ray crystallographic analysis of HALC3_919 with support from AK and AB. MAT performed pseudovirus entry assays. MAT and AA carried out flow cytometry and membrane fusions assays. NJC, HF and AS carried out *in vivo* studies. RB, DB and DV supervised the work and were responsible for funding acquisition. MAT, RJR, JL, DB and DV analyzed the data. MAT, RJR, DB and DV wrote the manuscript with input from all authors.

## Supplementary Tables

**Table S1.**
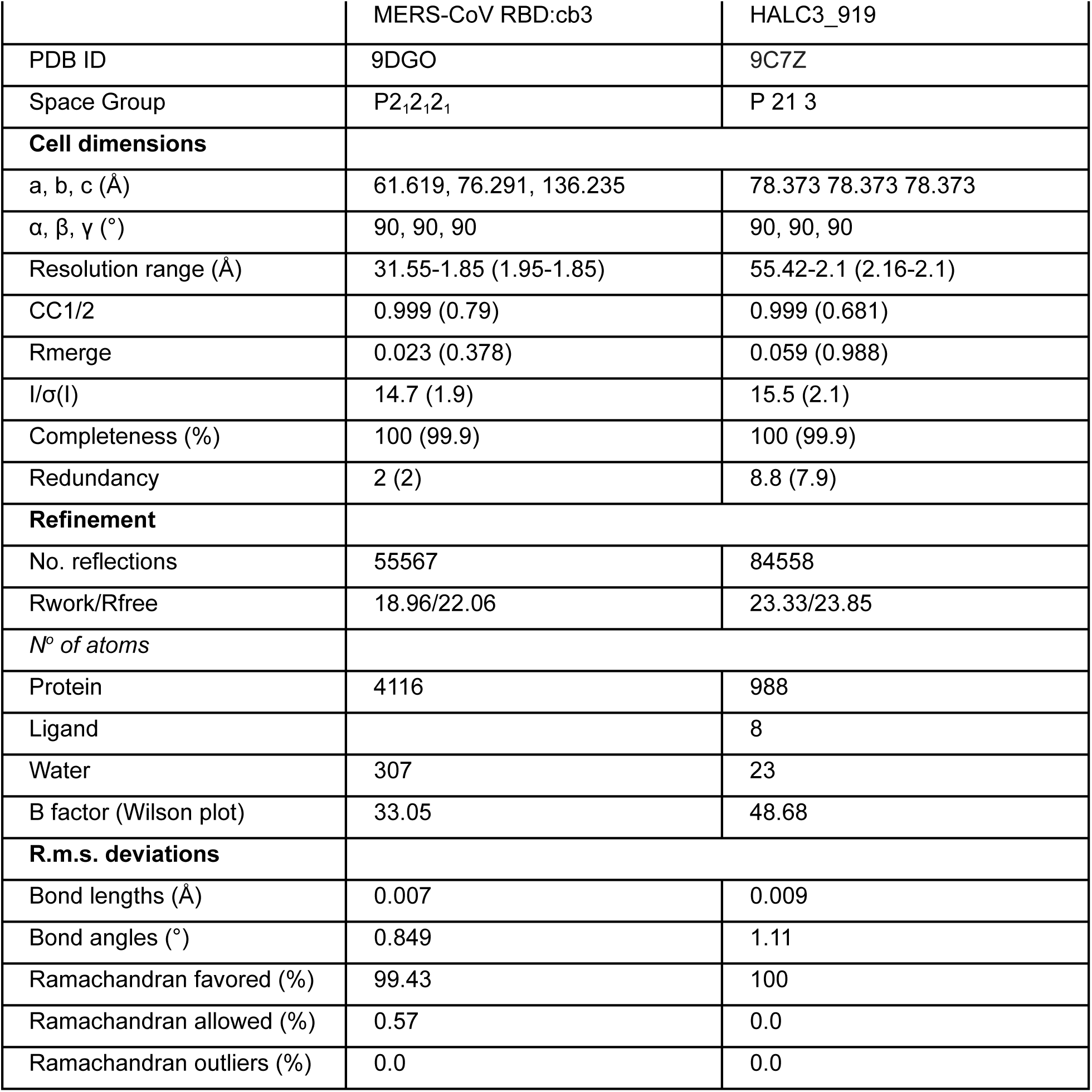
X-ray crystallography data collection and refinement statistics of the cb3-MERS-CoV RBD complex. - Data in parentheses are for the highest resolution shell - *R*_merge_ = ∑(∑|*I*_i_ − 〈I〉|/∑|*I*|), where the first ∑ is the sum over all reflections, and the second ∑ is the sum over all measurements of a given reflection, with *I*_i_ being the ith measurement of the intensity of the reflection and 〈*I*〉 the average intensity of that reflection. - *R*_work_/*R*_free_ = ∑(|*F*_o_| - 〈|*F*_c_|〉)/∑|*F*_o_|, where 〈|*F*_c_|〉 is the expectation of |*F*_c_| under the probability model used to define the likelihood function. The sum is overall reflections.

|  | MERS-CoV RBD:cb3 | HALC3_919 |
| --- | --- | --- |
| PDB ID | 9DGO | 9C7Z |
| Space Group | P2<sub>1</sub>2<sub>1</sub>2<sub>1</sub> | P 21 3 |
| **Cell dimensions** |  |  |
| a, b, c (Å) | 61.619, 76.291, 136.235 | 78.373 78.373 78.373 |
| α, β, γ (°) | 90, 90, 90 | 90, 90, 90 |
| Resolution range (Å) | 31.55-1.85 (1.95-1.85) | 55.42-2.1 (2.16-2.1) |
| CC1/2 | 0.999 (0.79) | 0.999 (0.681) |
| Rmerge | 0.023 (0.378) | 0.059 (0.988) |
| I/σ(I) | 14.7 (1.9) | 15.5 (2.1) |
| Completeness (%) | 100 (99.9) | 100 (99.9) |
| Redundancy | 2 (2) | 8.8 (7.9) |
| **Refinement** |  |  |
| No. reflections | 55567 | 84558 |
| Rwork/Rfree | 18.96/22.06 | 23.33/23.85 |
| *N° of atoms* |  |  |
| Protein | 4116 | 988 |
| Ligand |  | 8 |
| Water | 307 | 23 |
| B factor (Wilson plot) | 33.05 | 48.68 |
| **R.m.s. deviations** |  |  |
| Bond lengths (Å) | 0.007 | 0.009 |
| Bond angles (°) | 0.849 | 1.11 |
| Ramachandran favored (%) | 99.43 | 100 |
| Ramachandran allowed (%) | 0.57 | 0.0 |
| Ramachandran outliers (%) | 0.0 | 0.0 |

**Table S2.**
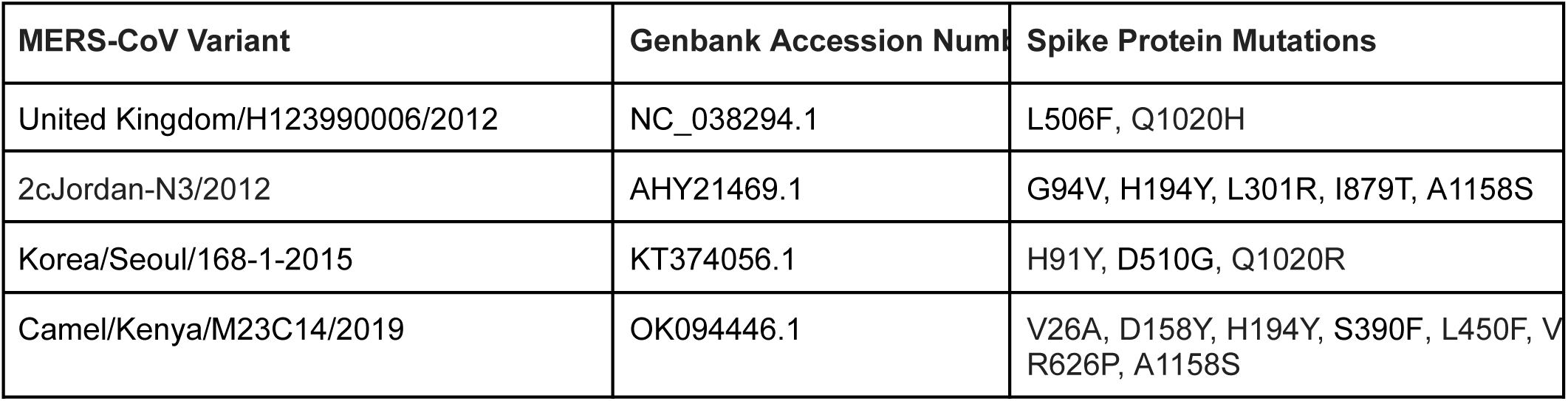
S glycoprotein mutations in MERS-CoV S variants used in this study relative to MERS-CoV S EMC/2012 (NC_019843.3). The mutations located at the receptor binding motif are highlighted in red.

| MERS-CoV Variant | Genbank Accession Numl | Spike Protein Mutations |
| --- | --- | --- |
| United Kingdom/H123990006/2012 | NC_038294.1 | L506F, Q1020H |
| 2cJordan-N3/2012 | AHY21469.1 | G94V, H194Y, L301R, I879T, A1158S |
| Korea/Seoul/168-1-2015 | KT374056.1 | H91Y, D510G, Q1020R |
| Camel/Kenya/M23C14/2019 | OK094446.1 | V26A, D158Y, H194Y, S390F, L450F, V<br>R626P, A1158S |

**Table S3.**
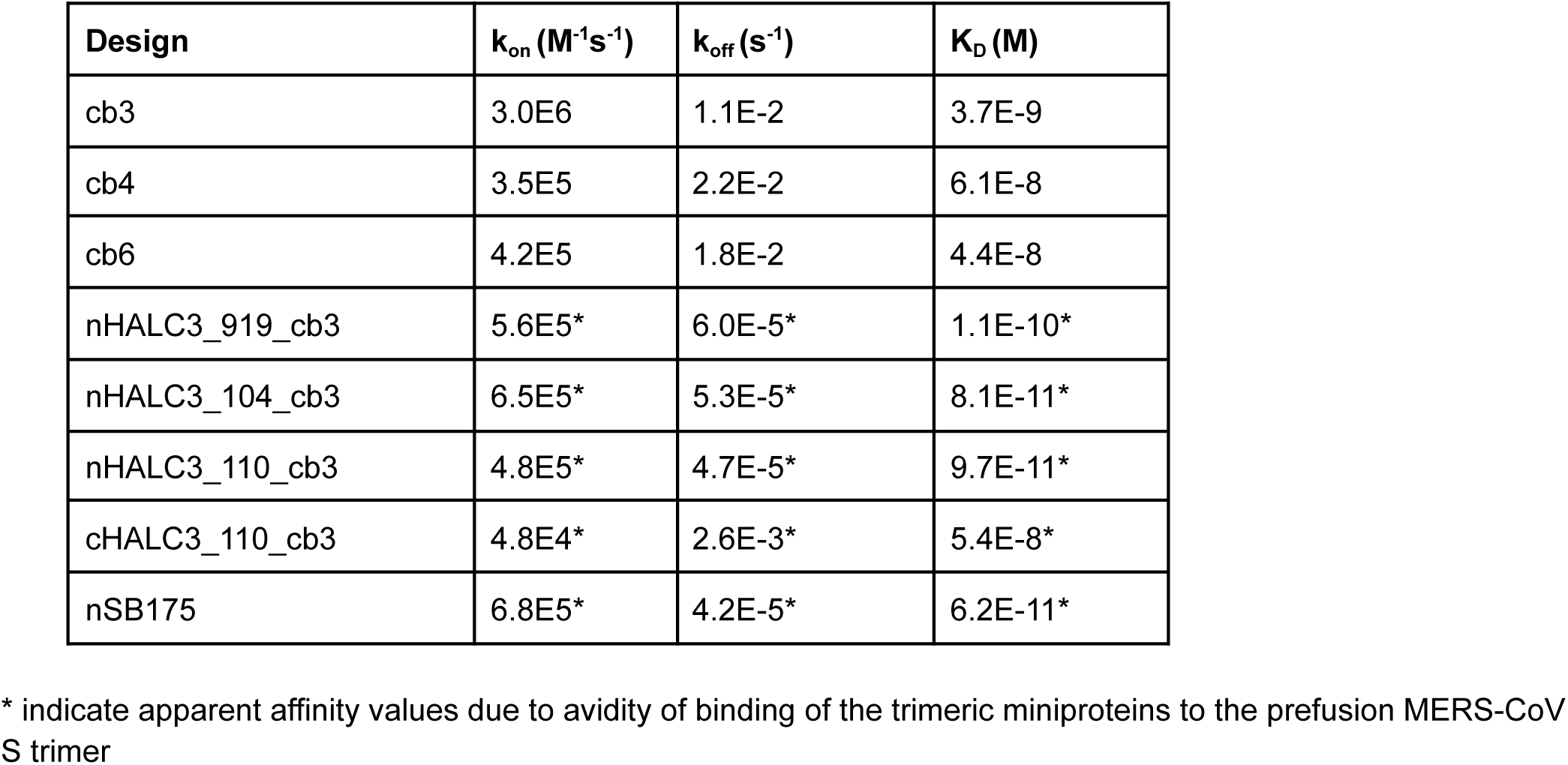
Summary of binding kinetics for monomeric and trimeric miniproteins. K_D_s were determined through global langmuir 1:1 model fitting.

**Table S4.**
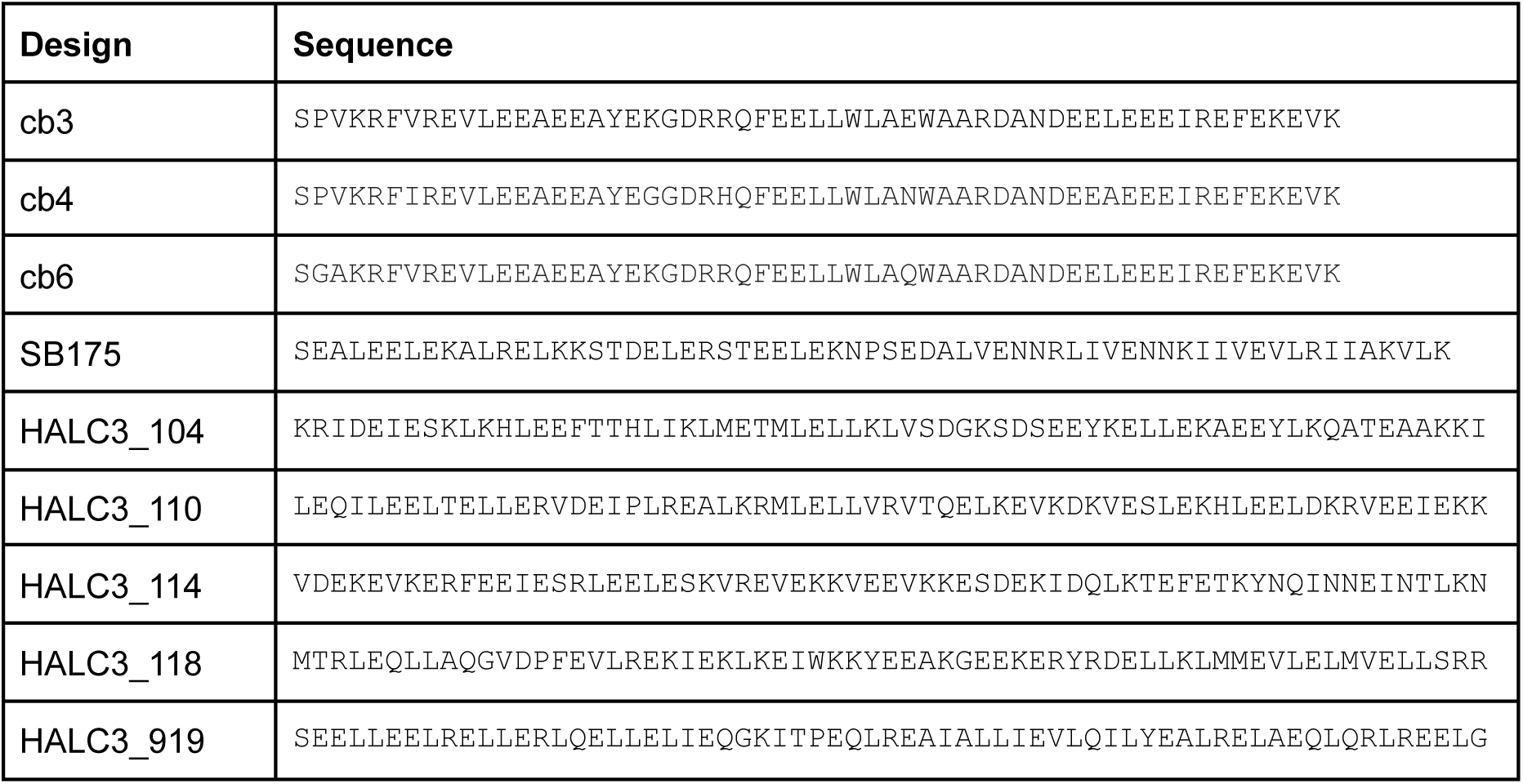
Designed sequences of miniproteins and homo-oligomer domains described in the text. Each trimer was tested as an N- and C-terminal fusion with respect to the miniprotein cb3. All constructs were expressed with MSG - design – GS - SNAC tag - 6x his, as described in methods.

**Table S5.**
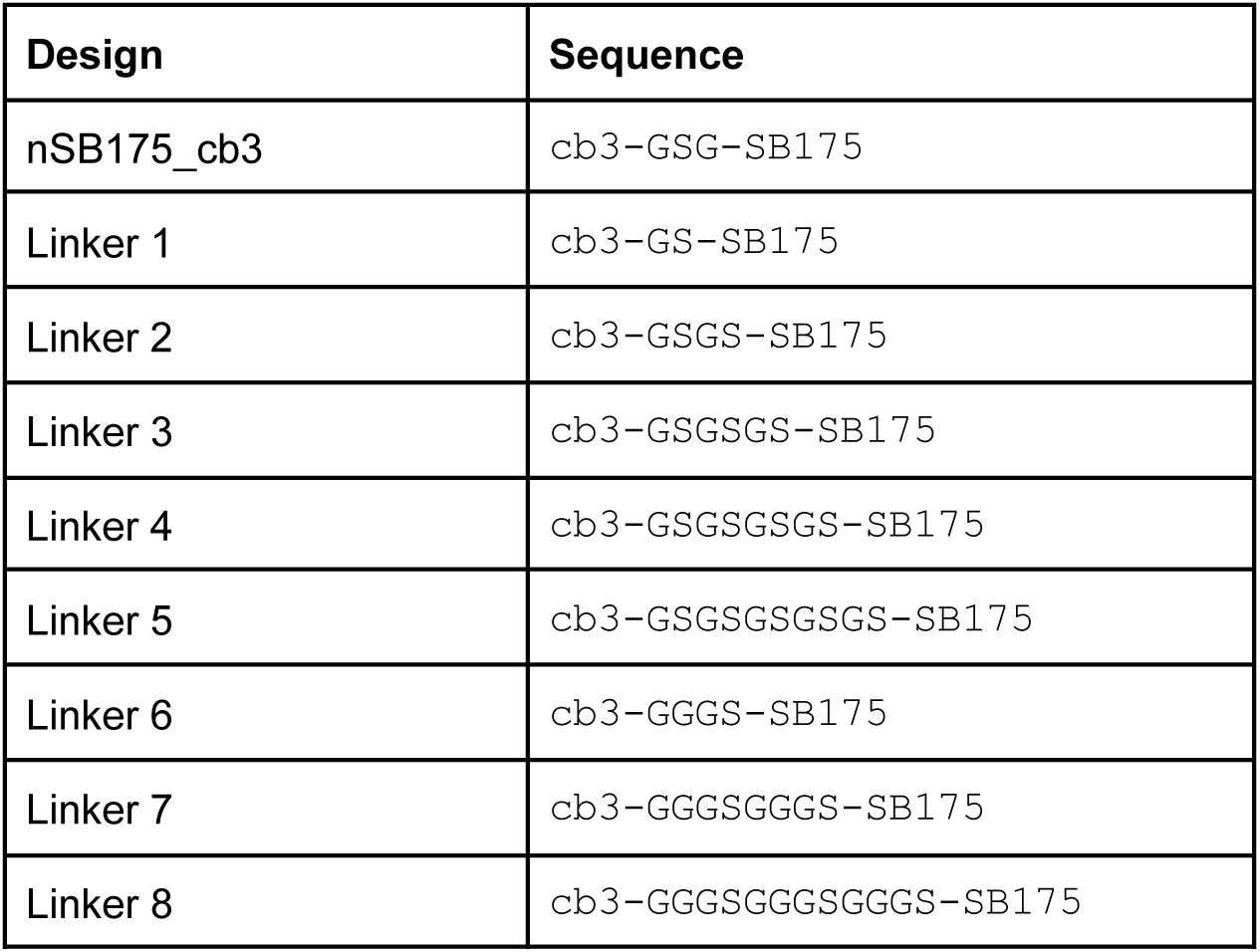
Sequences of the different linkers used between the trimerization domain nSB175 and the miniprotein cb3. The “n” refers to the position of the miniprotein cb3 relative to the trimerization domain. All constructs were expressed with MSG preceding the designed sequence and SNAC tag - 6x his at the C terminus, as described in methods.

**Table S6.**
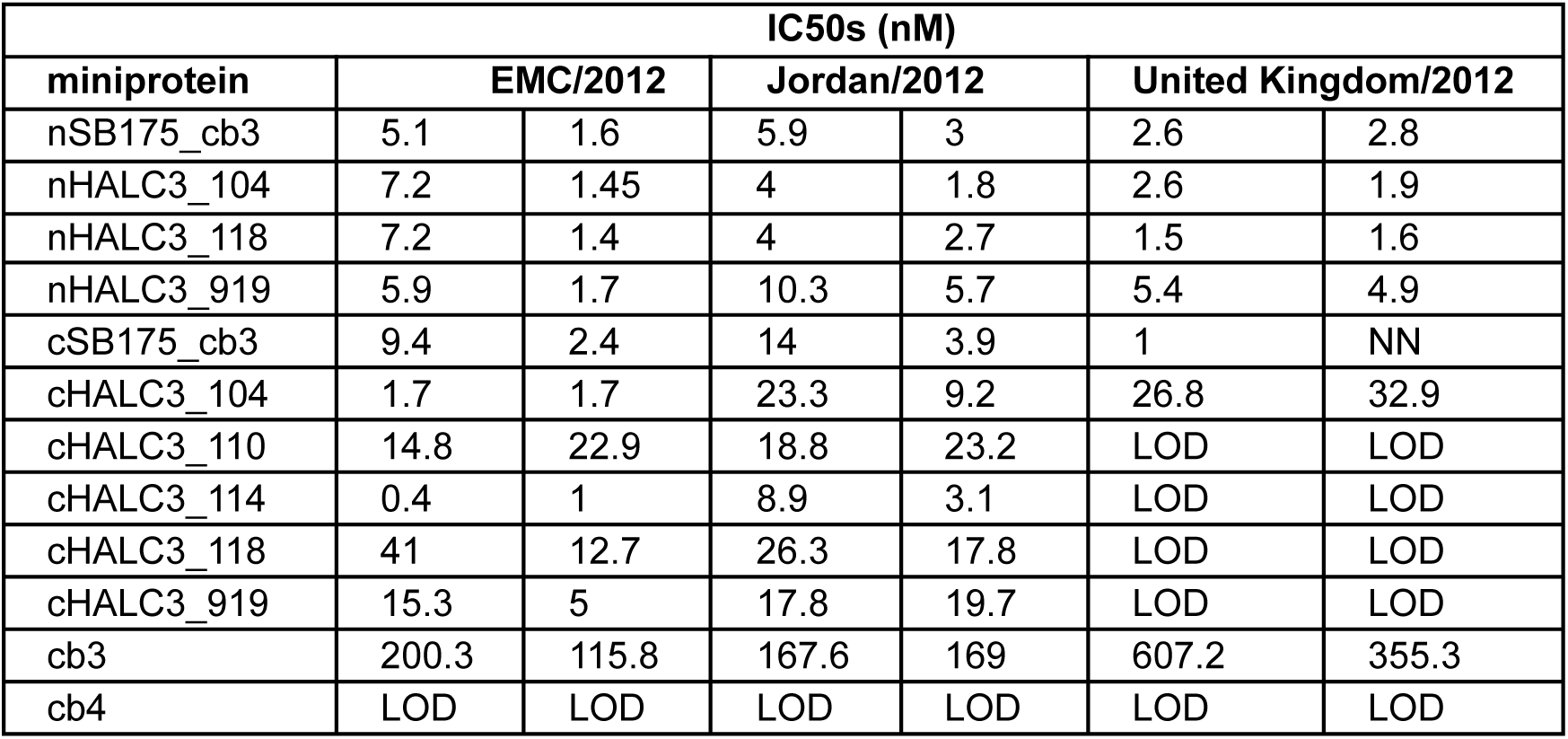
List of IC_50_ values obtained from the neutralization curves shown in Fig S3 expressed in nanomolar. Monomeric miniprotein cb3 was used as a reference to highlight the improved neutralization exhibited by the trimerization of cb3. Monomeric miniprotein cb4 was used as a negative control. The two IC_50_ values for each pseudotyped virus correspond to two distinct biological experiments performed with two batches of pseudovirus and one batch or miniprotein. The “n” and “c” indices refer to the position N- or C-terminus of the miniprotein cb3 relative to the indicated trimerization domain. NN: No Neutralization. Limit of detection (LOD) of miniproteins is between 5×10^2^-10^3^ nM (see Fig S3).

**Table S7.**
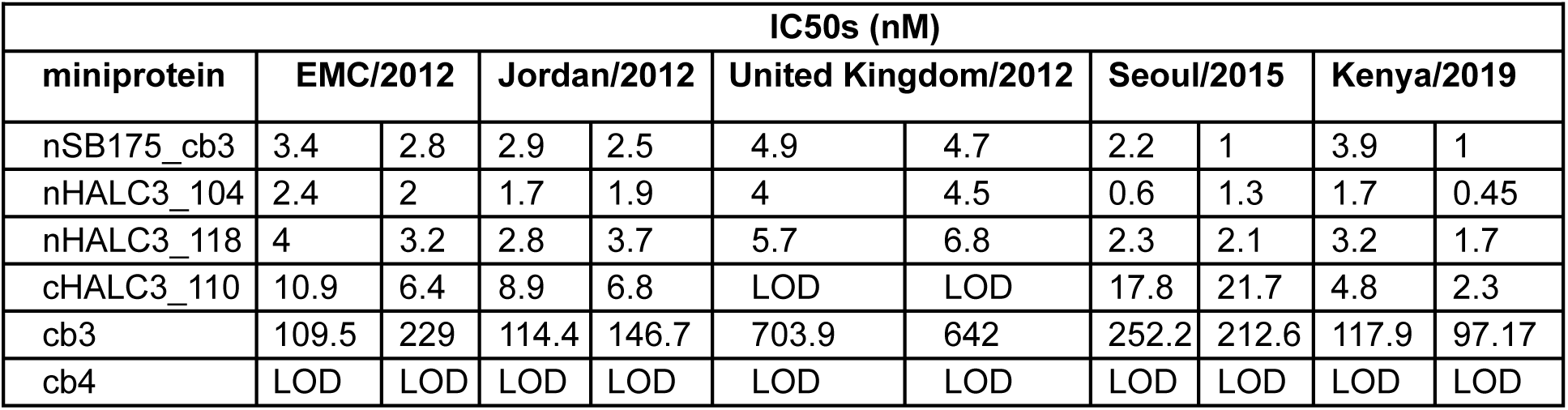
IC_50_ values obtained from the experiments shown in Fig S4 are expressed in nanomolar. Monomeric miniprotein cb3 was used as a reference and monomeric miniprotein cb4 was used as a negative control. The two IC_50_ values correspond to two different biological replicates using two batches of pseudovirus and one batch or miniprotein. The “n” and “c” indices refer to the position N- or C-terminus of miniprotein cb3 relative to the indicated trimerization domain. Limit of detection (LOD) for cb4 and cHALC3_110 is 10^3^ nM (see Fig S4).

**Table S8.** CryoEM data collection and refinement statistics.

| Data collection and processing | MERS-CoV S in complex with cb3-GSG-SB175, linker 1 (Global refinement, 2 RBDs engaged) | MERS-CoV S in complex with cb3-GSG-SB175, linker 1 (Global refinement, 3 RBDs engaged) | MERS-CoV S in complex with cb3-GGGSGGGS-SSB175, linker 7 (Global refinement) | MERS-CoV S in complex with cb3-GGGSGGGS-SSB175, linker 7 (Local refinement) | MERS-CoV S in complex with cb3-GGGSGGGS-SB175, linker 7 (Global refinement after focused classification, 3 RBDs engaged) |
| --- | --- | --- | --- | --- | --- |
| Magnification | 105,000 | 105,000 | 105,000 | 105,000 | 105,000 |
| Voltage (kV) | 300 | 300 | 300 | 300 | 300 |
| Electron exposure (e<sup>-</sup>/Å<sup>2</sup>) | 53.25 | 53.25 | 53.25 | 53.25 | 53.25 |
| Defocus range (-μm) | 0.8 - 1.7 | 0.8 - 1.7 | 0.8 - 1.7 | 0.8 - 1.7 | 0.8 - 1.7 |
| Pixel size (Å) | 0.829 | 0.829 | 0.829 | 0.829 | 0.829 |
| Symmetry imposed | C1 | C1 | C1 | C1 | C1 |
| Initial number of particles | 615,255 | 615,255 | 986,712 | 986,712 | 986,712 |
| Final number of particles | 271.037 | 79.019 | 538.654 | 169.944 | 63.523 |
| Map resolution (Å) | 2.9 | 3.3 | 2.6 | 3.2 | 3.0 |
| FSC threshold | 0.143 | 0.143 | 0.143 | 0.143 | 0.143 |
| Map sharpening B factor (Å<sup>2</sup>) | -12.13 | 19.11 | -46.60 | -66.35 | -21.03 |
| **Validation** |  |  |  |  |  |
| MolProbity score | N/A | N/A | N/A | 1.25 | N/A |
| Clash score | N/A | N/A | N/A | 4.75 | N/A |
| Poor rotamers (%) | N/A | N/A | N/A | 0.00 | N/A |
| **Ramachandran plot** |  |  |  |  |  |
| Favored (%) | N/A | N/A | N/A | 98.56 | N/A |
| Allowed (%) | N/A | N/A | N/A | 1.44 | N/A |
| Disallowed (%) | N/A | N/A | N/A | 0.0 | N/A |
| **Entry codes** | EMD:46947 | EMD:-46952 | EMD:46955 | PDB: 9DKK<br>EMDB: 46960 | EMD:46957 |

## Supplementary Figures

**Fig S1.**
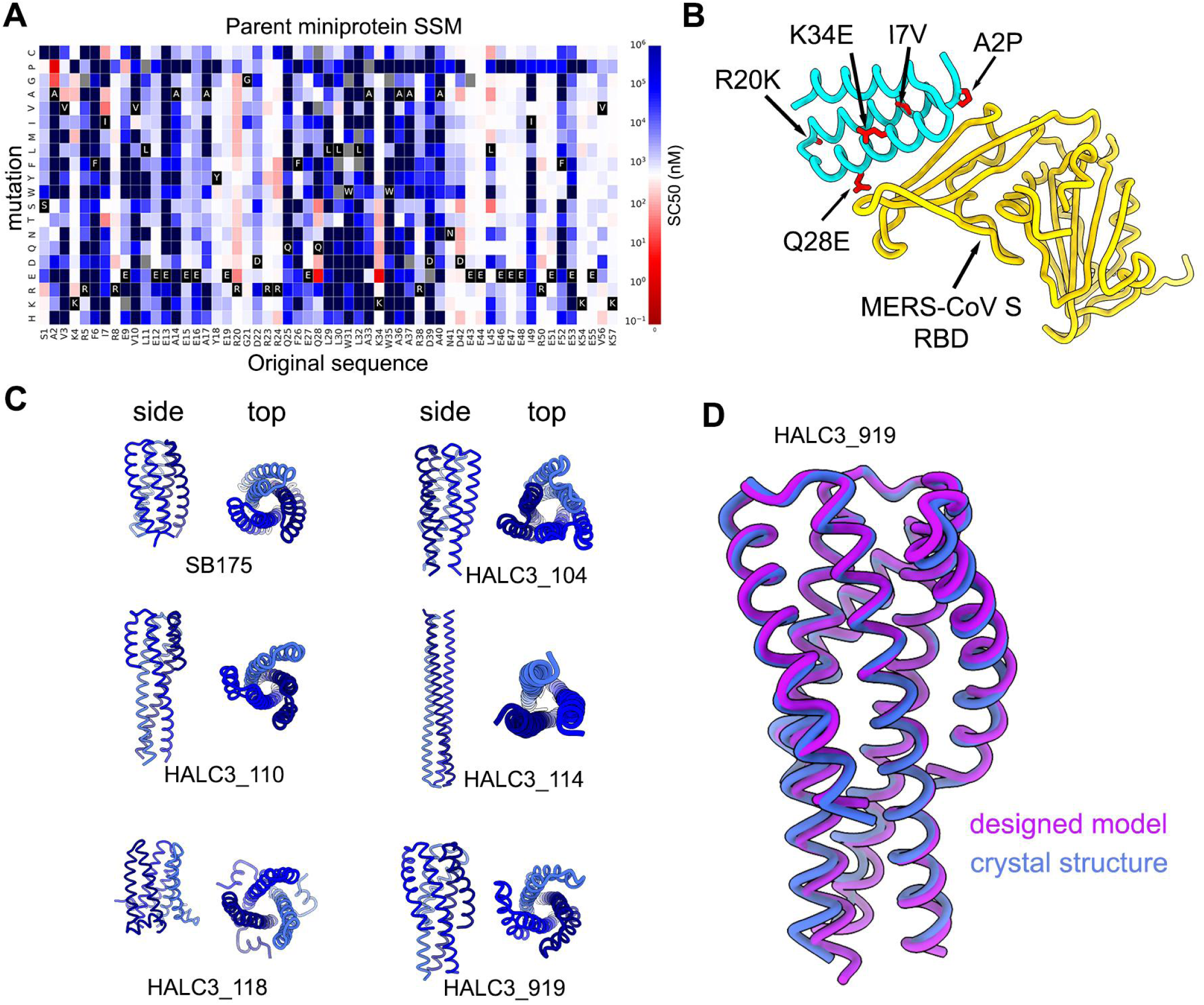
Optimization of MERS-CoV S RBD-directed miniproteins. **A.** Site saturation mutagenesis of the initial hit, designated parent miniprotein from which a combinatorial yeast surface display library was produced by selecting affinity enhancing point mutations. For each point mutation, the concentration that achieves 50 % of the saturating binding signal on yeast (SC50*(32)*) was calculated and plotted as enhancing affinity (red), reducing affinity (blue) or not affecting affinity (white) as compared to the SC50 of the parental design. The amino acid residue identity at each position of the parent miniprotein is colored black with white text, with all possible amino acid substitutions for that position following the y-axis. Gray indicates mutations that were not present in the library. **B.** Selected mutations from the parent miniprotein to yield the cb3 design are indicated on the ribbon diagram of the complex X-ray structure. **C**. Models of trimerization motifs fused N- or C-terminally to the cb3 miniprotein. **D**. Crystal structure of the HALC3_919 trimerization domain superposed to the design model, yielding an r.m.s.d of 1.52 Å across all Cα in the homotrimer.

**Fig S2.**
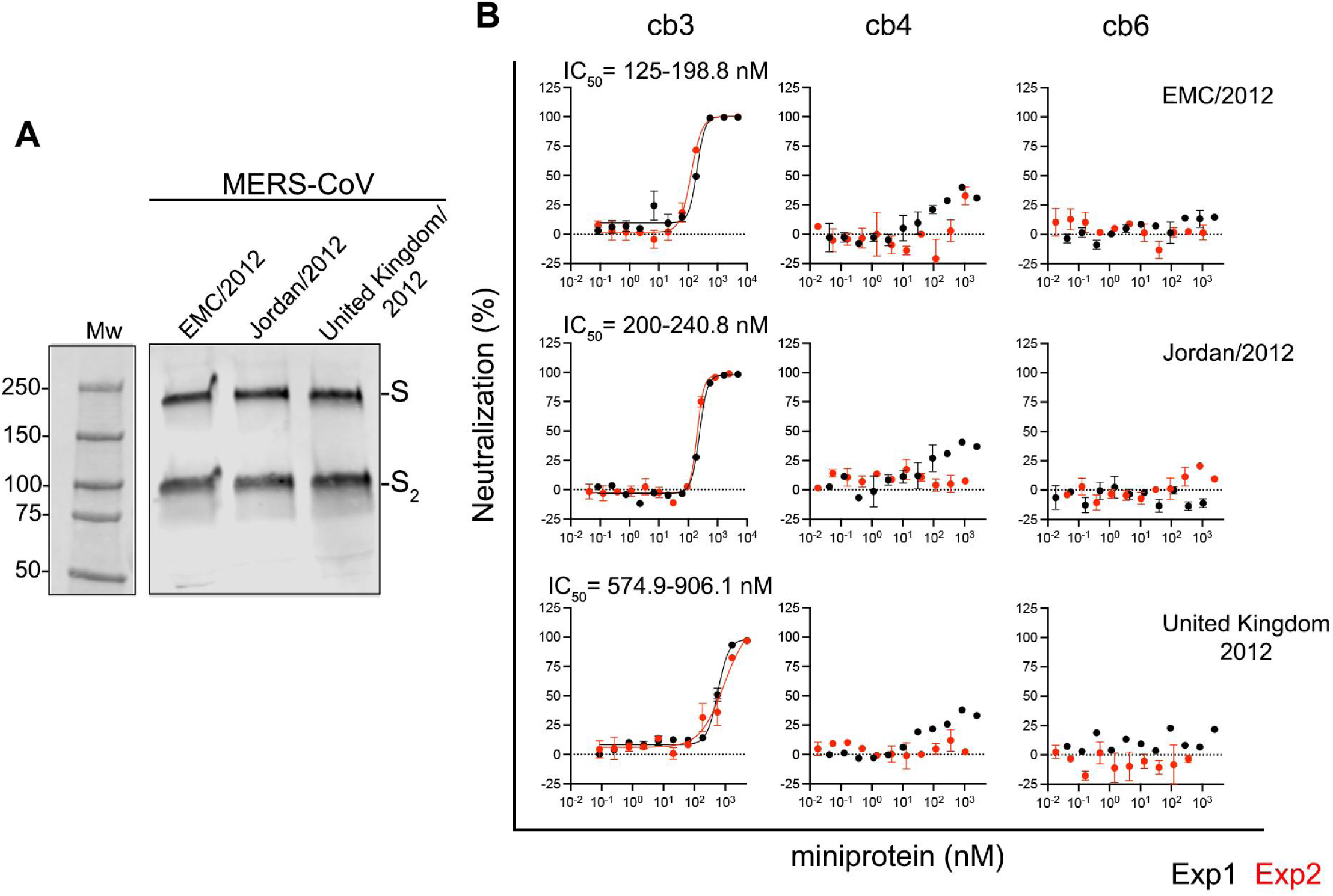
Inhibition of MERS-CoV S-mediated entry into VeroE6-TMPRSS2 cells by monomeric designed miniproteins. **A**, Western blot analysis of VSV pseudotyped particles harboring MERS-CoV EMC/2012, Jordan/2012 or United Kingdom/2012 S detected using the B6 stem-helix monoclonal antibody*(48)* as a primary antibody. Full-length S and S_2_ subunit bands are indicated on the right-hand side of the blot. **B**, Concentration-dependent inhibition of MERS-CoV S pseudovirus entry into VeroE6-TMPRSS2 cells for MERS-CoV S EMC/2012, Jordan/2012 and United Kingdom/2012. Exp 1 and Exp 2 correspond to two biological experiments performed with different preparations of pseudotyped viruses and one preparation of miniprotein. Error bars represent the standard error of the mean (SEM) of the technical duplicates. Fits are shown only when neutralization was detected.

**Fig S3.**
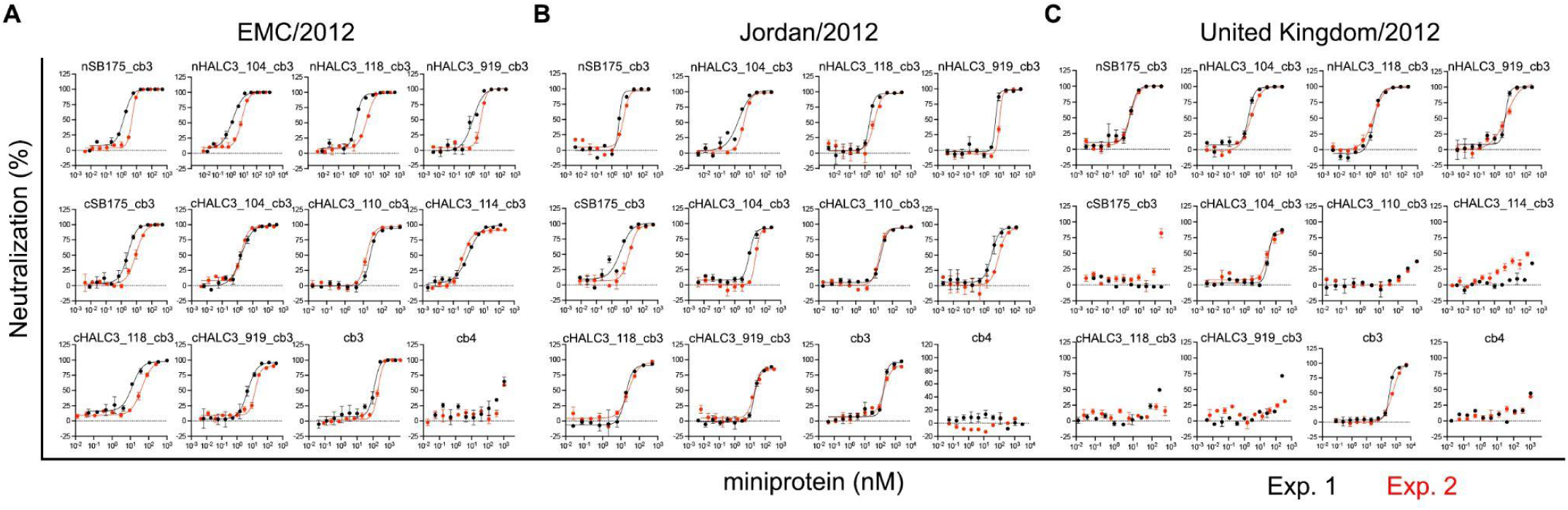
Inhibition of MERS-CoV S-mediated entry into VeroE6-TMPRSS2 cells by trimeric miniproteins. **A-C**, MERS-CoV EMC/2012 (**A**), Jordan/2012 (**B**) and United Kingdom/2012 (**C**) S VSV pseudovirus-mediated entry in the presence of various dilutions of the indicated miniproteins. Monomeric miniprotein cb3 was used as a reference and cb4 as negative control of neutralization. Exp 1 and Exp 2 correspond to two biological experiments performed with different preparations of pseudotyped viruses and one preparation of miniprotein. Error bars represent the standard error of the mean (SEM) of the technical duplicates. Fits are shown only when neutralization was detected.

**Fig S4.**
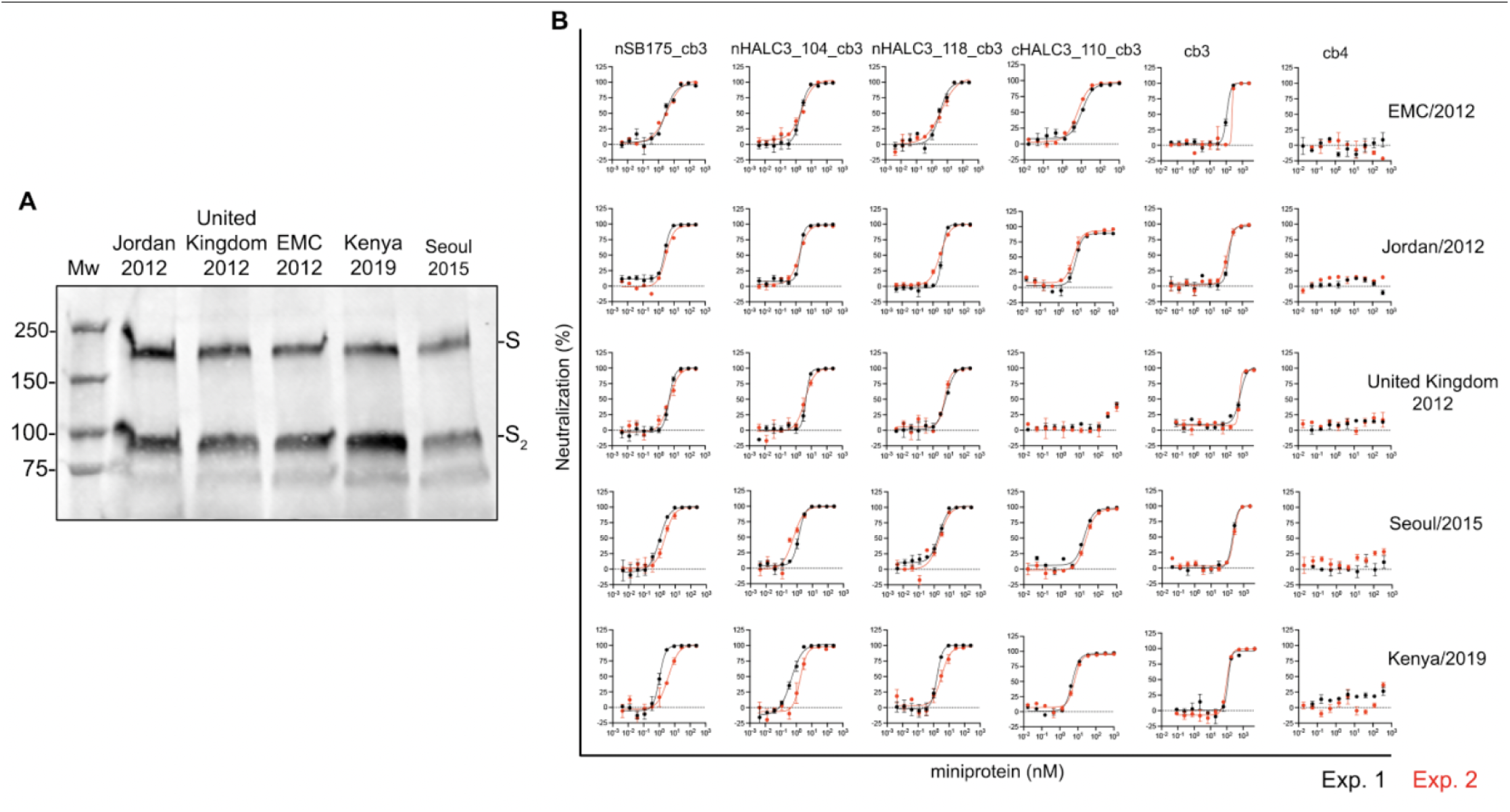
Inhibition of MERS-CoV S-mediated entry into VeroE6-TMPRSS2 cells by selected trimeric miniproteins. **A.** Western blot analysis of VSV pseudotyped particles harboring the indicated MERS-CoV S variants detected using the stem-helix monoclonal antibody B6*(48)* as a primary antibody Mw, molecular weight ladder. Full-length S and S_2_ subunit bands are indicated on the right-hand side of the blot. **B.** MERS-CoV EMC2012, Jordan/2012, United Kingdom/2012, Kenya/2019 and Seoul/2015 S VSV pseudovirus entry in the presence of various dilutions of the indicated miniproteins. Exp 1 and Exp 2 correspond to two biological experiments performed with different preparations of pseudotyped viruses and one preparation of miniprotein. Error bars represent the standard error of the mean (SEM) of the technical duplicates. Fits are shown only when neutralization was detected.

**Fig S5.**
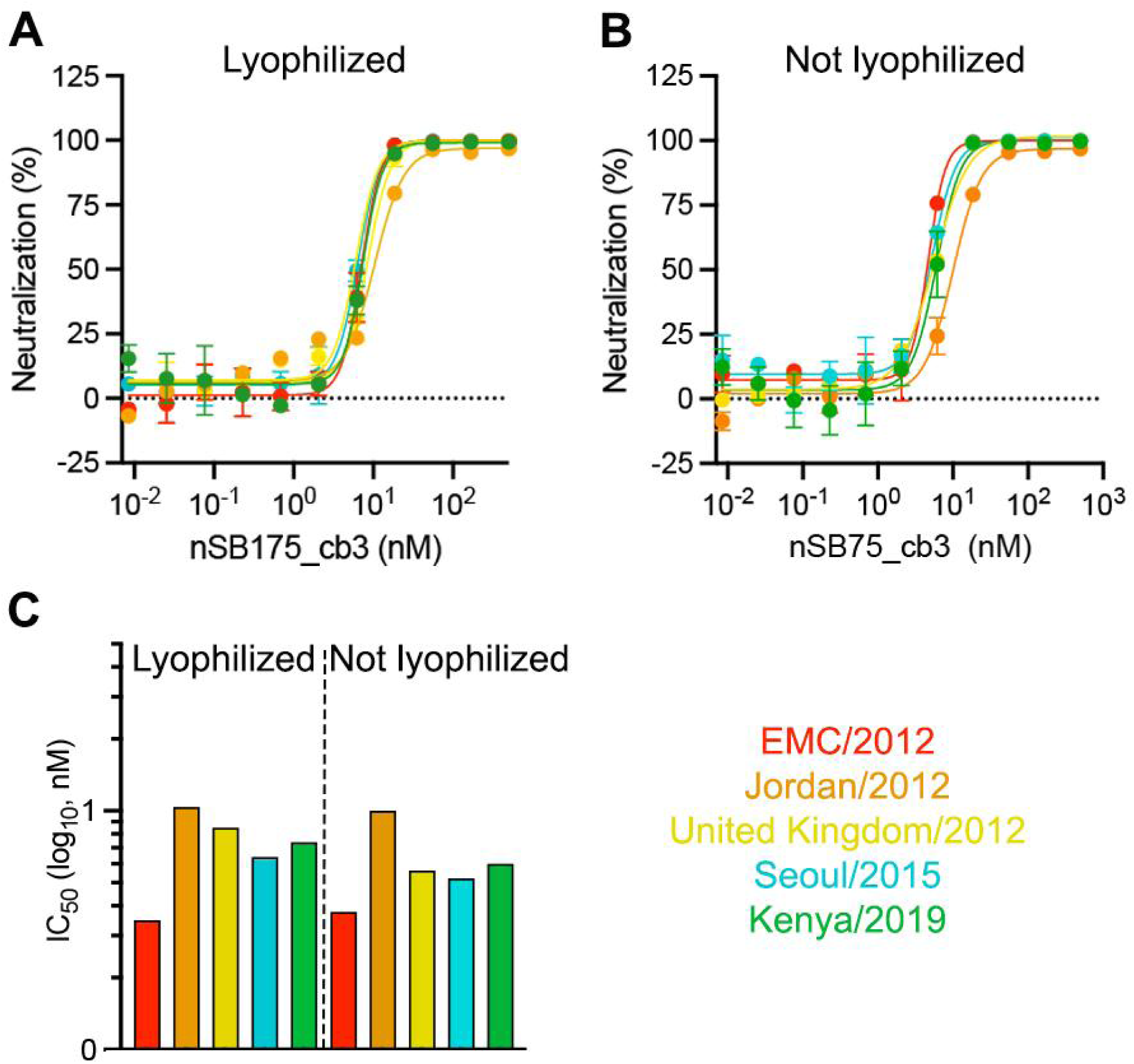
Retention of neutralizing activity upon miniprotein nSB175_cb3 (cb3-GSG-SB175) lyophilization. MERS-CoV EMC/2012, Jordan/2012, United Kingdom/2012, Kenya/2019 and Seoul/2015 S pseudovirus entry in the presence of various dilutions of nSB175_cb3 lyophilized and reconstituted (**A**) or not lyophilized (**B**). A single biological experiment with technical duplicates is shown. Error bars represent the standard error of the mean (SEM) of the technical duplicates. **C**. IC_50_ values, expressed in nanomolar, obtained from the experiment shown in panels A and B.

**Fig S6.**
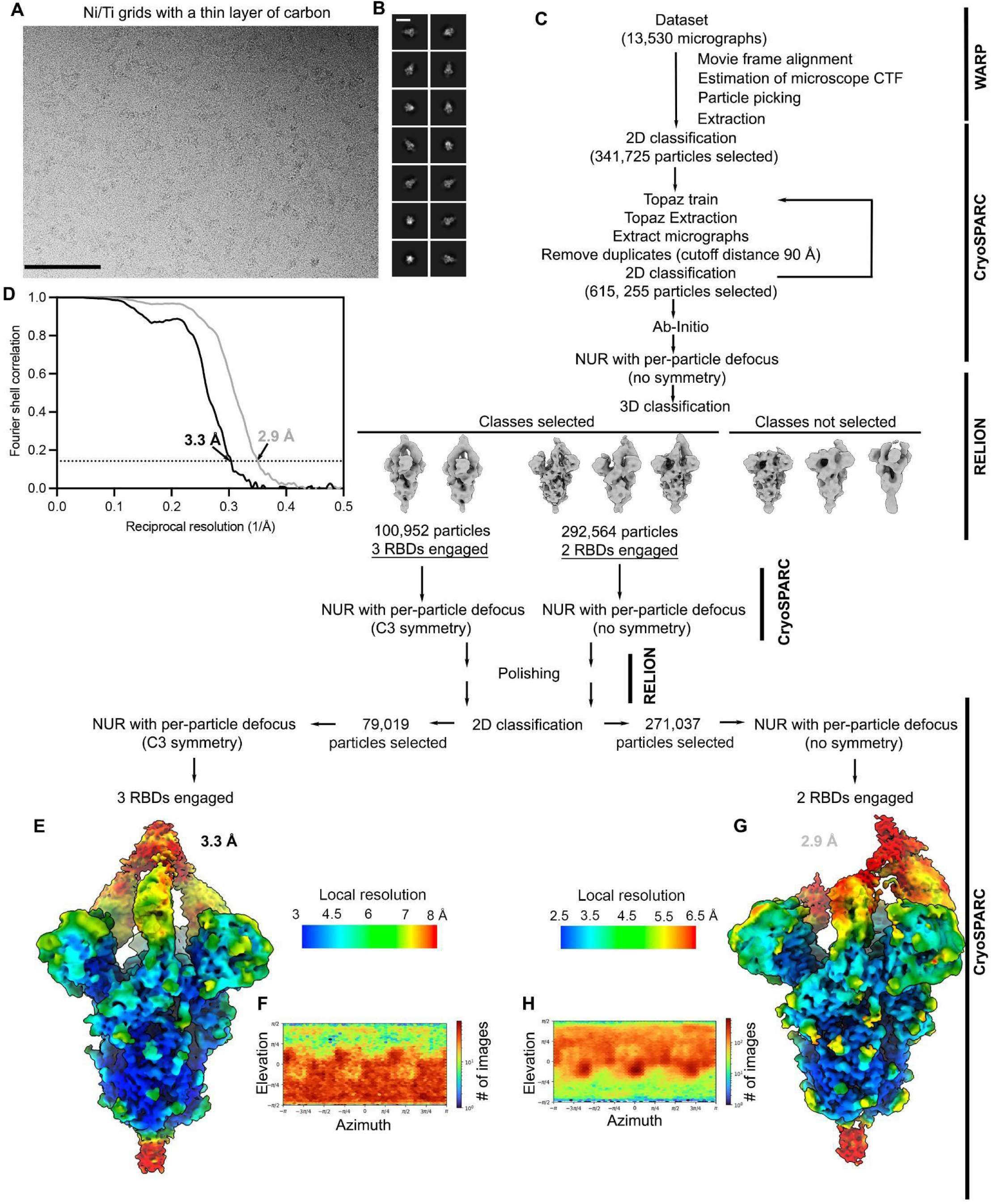
CryoEM data processing and validation of the structure of MERS-CoV S in prefusion conformation in complex with nSB175_cb3 (cb3-GSG-SB175). **A.** Representative electron micrograph. **B.** 2D class averages. Scale bar of the micrograph and the 2D class averages, 100 nm and 100 Å, respectively. **C.** Cryo-EM data processing flowchart. CTF: contrast transfer function. NUR: non uniform refinement. **D.** Gold-standard Fourier shell correlation curves for the global maps with three and two RBDs engaged are shown in black and gray, respectively. The 0.143 cutoff is indicated by a horizontal dotted black line. **E.** Unsharpened map corresponding to prefusion MERS-CoV S in complex with three nSB175_cb3 colored by local resolution. **F.** Angular distribution plot with all the particles contributing to the map in D. **G.** Unsharpened map corresponding to the prefusion MERS-CoV S in complex with two nSB175_cb3 colored by local resolution. **H.** Angular distribution plot with all the particles contributing to the map in G.

**Fig S7.**
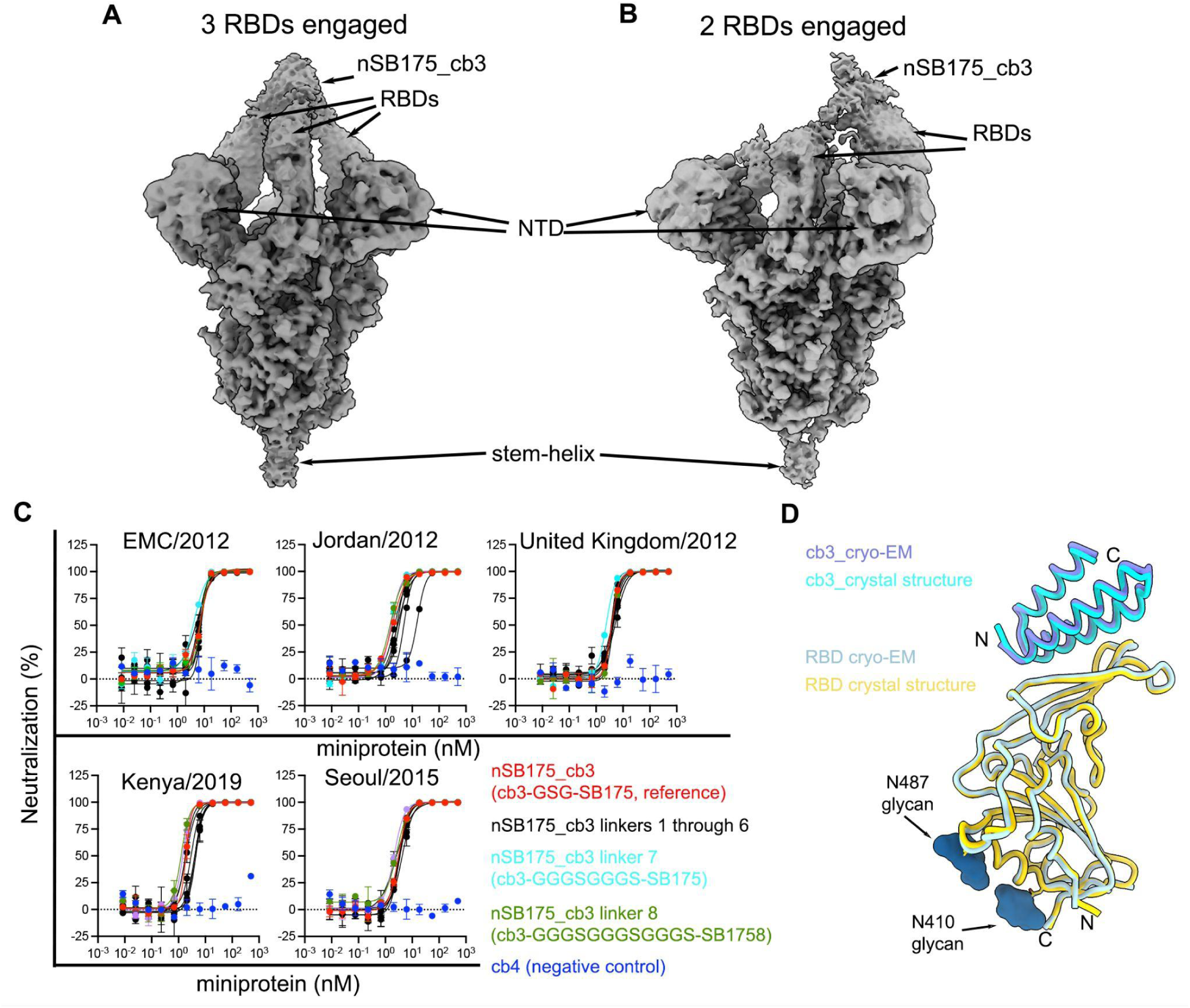
Optimization of the linker length between the miniprotein binding domain cb3 and the trimerization domain SB175. **A-B**. Unsharpened cryoEM maps corresponding to MERS-CoV S bound to three (**A**) or two (**B**) cb3 modules simultaneously from the nSB175_cb3 homotrimeric miniprotein. **C.** MERS-CoV EMC/2012, Jordan/2012, United Kingdom/2012, Kenya/2019 and Seoul/2015 S VSV pseudovirus entry into cells in the presence of various dilutions of nSB175_cb3 with different linkers sizes between cb3 and the trimerization domain. Miniprotein cb4 was used as a negative control. A single biological experiment with two technical replicates is shown. Error bars represent the standard error of the mean (SEM) of the technical duplicates. **D.** Structural overlay between the cryo-EM structure of the MERS-CoV S RBD in complex with nSB175_cb3 linker 7 and the X-ray structure of the MERS-CoV S RBD in complex with a single cb3.

**Fig S8.**
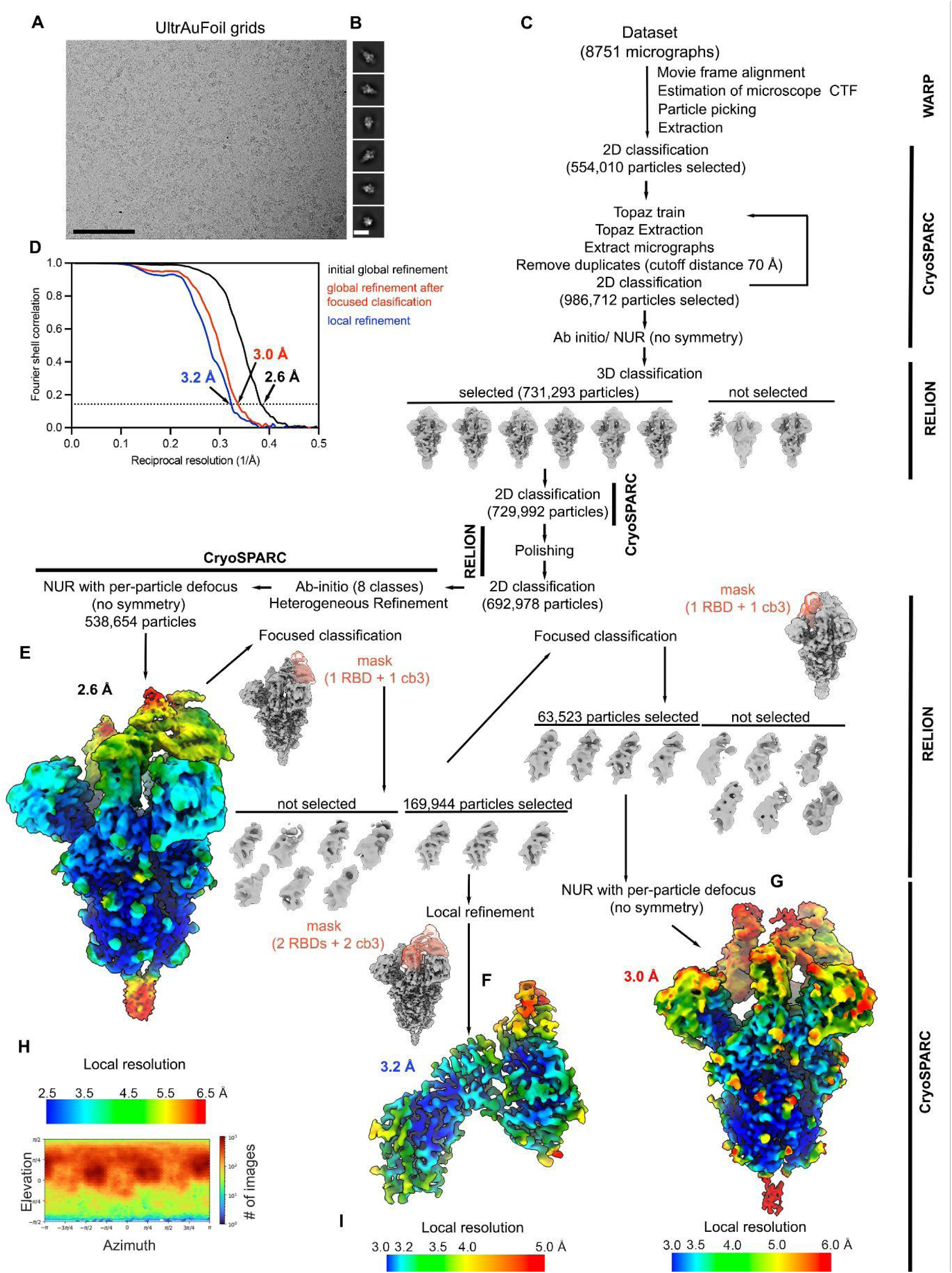
CryoEM data processing and validation of the structure of prefusion MERS-CoV S in complex with nSB175_cb3 linker 7 (cb3_GGGSGGGS_SB175). **A.** Representative electron micrograph. **B.** 2D class averages. Scale bar of the micrograph and the 2D class averages, 100 nm and 100 Å, respectively. **C.** Cryo-EM data processing flowchart. CTF: contrast transfer function. NUR: non uniform refinement. **D**. Gold-standard Fourier shell correlation curves for the global maps (black and red) and locally refined map (blue). The 0.143 cutoff is indicated by a horizontal dotted black line. **E.** Unsharpened map corresponding to the 3D reconstruction of MERS-CoV S (in prefusion conformation) in complex with nSB175_cb3 linker 7 colored by resolution. **F.** Local refined sharpened map corresponding to two MERS-CoV S RBDs engaging two cb3 from the nSB175_cb3 linker 7 miniprotein colored by resolution. **G.** Global unsharpened map for the MERS-CoV-S in complex with nSB175_cb3 linker 7 miniprotein obtained after focused classification and colored by resolution. **H, I.** Angular distribution plots corresponding to the maps shown directly above.

**Fig S9.**
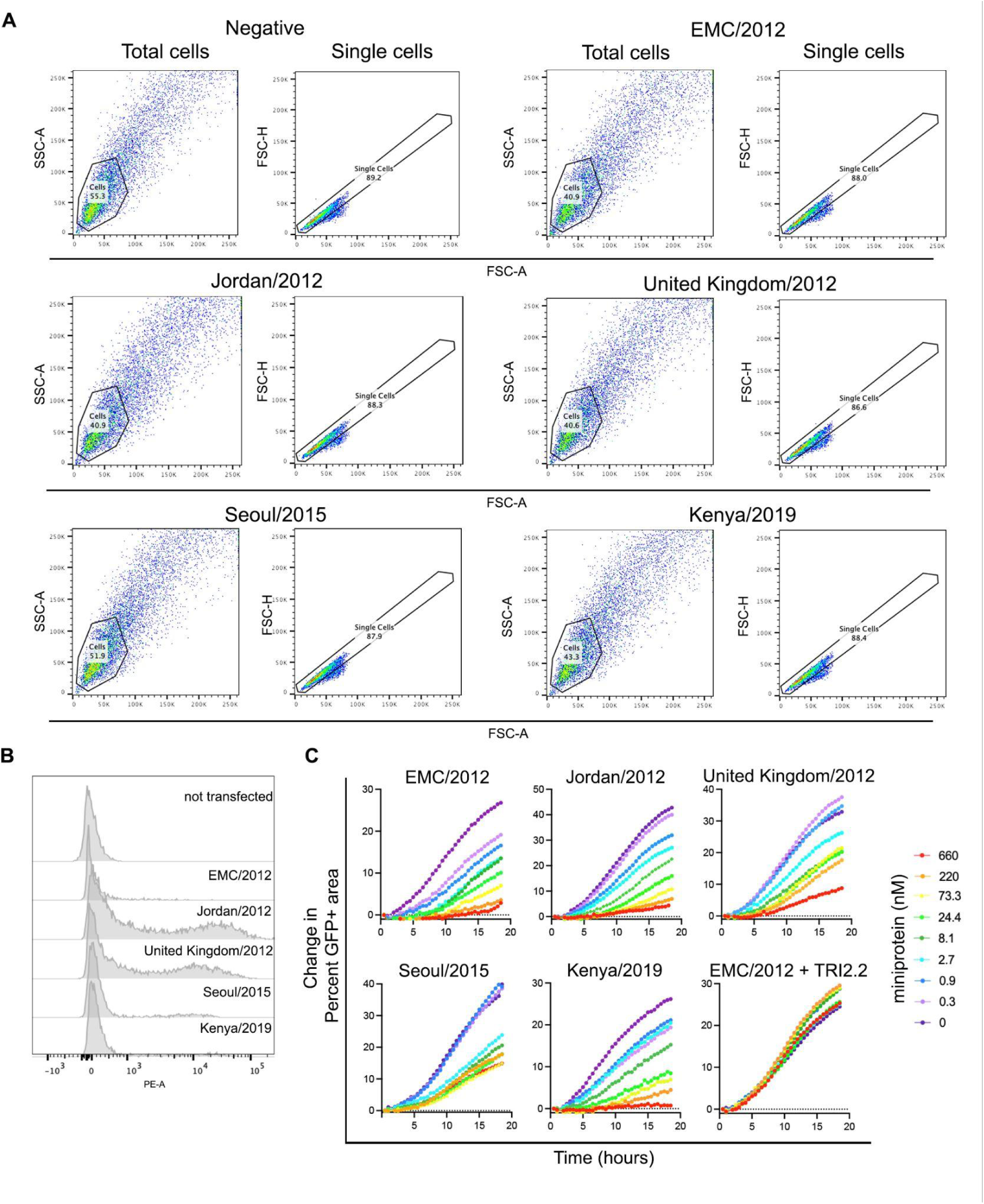
nSB175_cb3 (cb3-GSG-SB175) inhibits MERS-CoV S-mediated membrane fusion. **A.** Gating strategy for flow cytometry analysis of cell-surface expression of MERS-CoV S proteins from EMC/2012, Jordan/2012, United Kingdom/2012, Kenya/2019 and Seoul/2015 strains expressed at the surface of BHK-21-GFP_1-10_ cells. Representative gating to exclude cell debris and dead cells (SSC-A/FSC-A, total cells) and to select single cells (SSC-H/FSC-A, single cells) are shown. **B.** Quantification by flow cytometry of the different MERS-CoV S surface expressed on BHK-21-GFP_1-10_ using the stem helix antibody B6*(48)*. The y-axis is present as a modal scale scaled to maximum singleton events for that plot. **C**. Kinetics of cell-cell fusion (expressed as a change in percent of GFP^+^ area) promoted by the different MERS-CoV S glycoproteins over an 18 h time course experiment using a split GFP system and in the presence or absence of different concentrations (expressed in nM) of cb3-GSG-SB175 or TRI2.2. Data represent one experiment out of two biological replicates.

**Fig S10.**
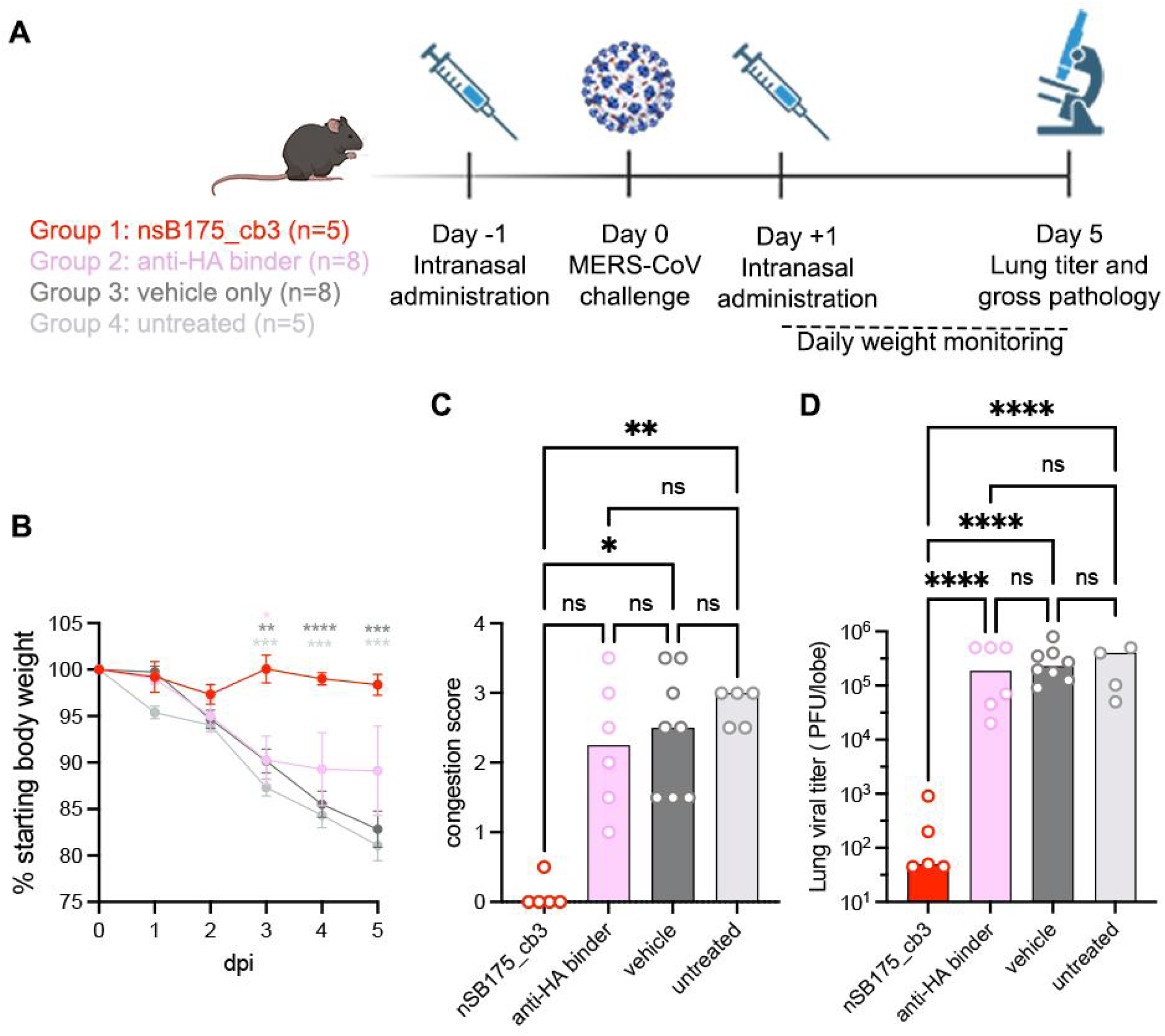
Pre- and post-administration of nSB175_cb3 (cb3-GSG-SB175) protects mice from MERS-CoV challenge. **A.** Trimeric fusion nSB175_cb3 (cb3-GSG-SB175), influenza hemagglutinin miniproteins (anti-HA), or vehicle alone were administered to mice C57BL/6 J 288/330 at day −1 and day +1. An untreated group was also included. Mice were challenged with MERS-CoV-m34c5 at day 0 and monitored for 5 days for body weight (**B**) or analyzed at day 5 for congestion score (**C**) and viral titer in lungs (**D**). Group comparisons for body weight and viral titers were assessed with the two-way ANOVA: Tukey’s test. For the congestion score, comparisons among groups were assessed with the Kruskal-Wallis test; ns, not significant; *P < 0.05, **P < 0.01, ****P < 0.0001.

## Materials and Methods

### Cell lines

Cell lines used in this study were obtained from ATCC (HEK293T), JCRB-Cell Bank (VeroE6-TMPRSS2), ThermoFisher Scientific (Expi293F™ cells), or generated in the lab (BHK-21-GFP_1-10_ or VeroE6-TMPRSS2-GFP_11_). Cells were cultivated at 37°C, in an atmosphere of 5 % CO_2_ and with 130 rpm of agitation for suspension cells. None of the cell lines used were routinely tested for mycoplasma contamination.

### Plasmids

All genes were synthesized by GenScript, codon optimized for expression in mammalian cells and in frame with a Kozak’s sequence to direct translation, except for the plasmid encoding membrane-anchored MERS-CoV EMC/2012 S, which was kindly provided by Whittaker lab. Genes encoding MERS-CoV RBD and MERS-CoV-2P Spike (S) ectodomain were cloned into pCMVR and pcDNA3.1 (+), respectively, with the signal peptide derived from the µ phosphatase: MGILPSPGMPALLSLVSLLSVLLMGCVAETGT. MERS-CoV RBD gene was fused at the c-terminus to an octahistidine tag while MERS-2P S protein was c-terminally fused to a foldon trimerization motif followed by a hexahistidine tag for affinity purification. Gene encoding human DPP4 was cloned in pDNA3.1 (+) with the signal peptide from the µ phosphatase and c-terminally fused to thrombin cleavage site followed by an FC-tag used for purification. Genes encoding MERS-CoV S full-length strains, Jordan/2012 (residues 1 to 1353), United Kingdom/2012 (residues 1-1353), Seoul/2015 (residues 1-1353) and Kenya/2019 (residues 1-1353) strains were untagged and with the native signal peptide.

Monomeric cb3, cb4 and cb6 were ordered as eblock fragments and inserted into the LM0627 plasmid (Aggene 191551) flanked by the MSG residues at the N-terminal and by a SNAC*(49)* and his-tag at the C-terminal. We used a BsaI golden gate assembly (NEB) in a 1 μL mixture reaction containing: 0.1 μL water, 0.1 μL T4 ligase buffer, 0.375 μL eblock fragment at 4 ng/µL, 0.06 μL of BsaI-HFv2, 0.1 μL T4 ligase, 0.275 μL of LM0627 vector at 50 ng/µL. Reactions were incubated for 30 minutes at 37°C before transformation of BL21 competent E. *coli* (NEB) cells. Plasmid sequences were confirmed through sanger sequences of the insert using T7 and T7-term primers from Azenta.

Trimerization domains were ordered as eblock fragments (IDT) with a ccdb selective marker (flanked by BsaI cut sites) either at the 5’ or 3’ of the trimerization domain and then inserted into a pET29b+ vector using PaqCI (NEB). Briefly, 2:1 molar ratio of trimerization domain:plasmid were ligated at 37°C for 20 minutes in a 10 µl mixture reaction containing: 1 μL T4 ligase buffer (NEB), 0.5 μL PaqCI (NEB), 0.5 μL T4 DNA ligase (NEB) and 0.25 μL PaqCI activator (NEB). Ligation reactions were then transformed into NEB-stable competent E. *coli* cells. After verifying the sequences of these plasmids, an eblock fragment encoding E. *coli* codon optimized cb3 was inserted into the pET29b+ containing the trimerization domains with the same golden gate assembly reaction described above.

### MERS-CoV RBD movalent and multivalent miniproteins design

The design of cb3 followed the same procedure as previous attempts to design binders against arbitrary targets of interest*(25, 32)*. First, a library of 30,000 helical bundles were docked onto the DPP4 interface focusing on W553, L506, V555 and Y540 on the MERS-CoV RBD from PDB 4KR0*(15)* using Rifgen-Patchdock-Rifdock *(31, 32, 50)*. The exact Rifdock settings used are provided in the supplementary rifdock.flag file. Rifdock outputs were then sorted using the predicted metrics after a minimal sequence design procedure with the best scoring designs from this short FastDesign implementation taken forward to full sequence design, a process described in depth previously*(32)*. After full sequence design for a subset of the best scoring docks, these designs were ranked by contact molecular surface, rosetta ΔΔg and contact molecular surface to critical hydrophobic residues. Interface secondary structural elements were extracted from the best designs, clustered, and ranked by ΔΔG to provide 3,000 *de novo* interface motifs. The 30,000 helical bundles were superimposed on these interface motifs to generate more docks which were subjected to the same design and ranking process as the Rifdock outputs. These motif-grafted outputs were combined with the Rifdock outputs and the best designs from the union ordered.

SB175 trimerization domain is a Rosetta-based design previously described*(24)*. All HALC3 trimerization domains were developed using AlphaFold2 hallucination*(37)*, except for HALC3_919 which is a symmetric homotrimer composed of three helical subunits of 65 amino acids. The protein backbone of HALC3_919 was generated using symmetric AlphaFold2 Markov Chain Monte Carlo hallucination. The sequence was generated using the Rosetta Fast Design function using a position specific scoring matrix generated from 9-mer fragments extracted from the PDB*(51, 52)*, and layer design limiting amino acids on surface exposed loops and strands to the following amino acids: DEGHKNPRQST.

### Recombinant protein production

MERS-CoV-RBD and MERS-CoV-2P S were expressed in Expi293F cells at 37°C and 8 % CO_2_. 100 ml of cells were transfected with 640 μL of Expifectamine and 100 µg of the respective plasmids, following the manufacturer’s indications. Four days post-transfection, supernatants were clarified by centrifugation at 3500 rpm for 15 minutes, supplemented with 300 mM NaCl and 25 mM sodium phosphate pH 7.4, bound to HisPur Cobalt Resin previously equilibrated with binding buffer (25 mM sodium phosphate pH 7.4 and 300 mM NaCl), and mixed on an end-over-end rotator at 4°C for one hour. Recombinant proteins were eluted using 25 mM sodium phosphate pH 7.4, 300 mM NaCl, and 500 mM imidazole. Proteins were further purified by size-exclusion chromatography using a Superdex 75 Increase 10/300 GL column pre-equilibrated in 25 mM Tris-HCl pH 8.0, 150 mM NaCl. Ectodomain of human DPP4 fused to an FC-tag was expressed following a similar protocol. After 4 days of expression, supernatants were clarified and passed through a HiTrap Protein A HP column (Cytiva) previously equilibrated in 20 mM sodium phosphate pH 8.0. Proteins were eluted using 0.1 M citric acid pH 3.0 in individual tubes (1 ml final volume per fraction) containing 200 μL of 1 M Tris-HCl pH 9.0 to immediately neutralize the low pH needed for elution. Fractions containing purified desired proteins were pooled and buffer exchanged to 25 mM Tris-HCl pH 8.0, 150 mM NaCl.

Miniprotein monomers and trimers were expressed from glycerol stocks inoculated into 50 ml of autoinduction media and grown at 37°C for 18 h. Cells were harvested via centrifugation at 4000 x g for 5 minutes followed by resuspension in 5 ml lysis buffer (20 mM Tris-HCl pH 8.0, 300 mM NaCl, 25 mM imidazole, 1 mM PMSF, 0.1 mg/ml lysozyme, 10 μg/ml DNAse I). Cells were lysed by sonication and then centrifuged at 13000 x g for 10 minutes to clarify the lysates before loading on to 0.5 mL Ni-NTA resin (Thermo Fisher). The resin was washed with 3 x 5 column volumes (CVs) of 20 mM Tris-HCl pH 8.0, 300 mM NaCl, 25 mM imidazole and then eluted in 20 mM Tris-HCl pH 8.0, 300 mM NaCl, 500 mM imidazole. Proteins were further purified via size exclusion chromatography into TBS (20 mM Tris-HCl pH 7.4, 150 mM NaCl) on a Superdex 75 10-300 GL Increase column for monomers and Superdex 200 10-300 GL Increase column for trimers.

For neutralization and *in vivo* studies, the samples were prepared with reduced endotoxin through the addition of four detergent washes while bound to the Ni-NTA resin using TBS with 0.75% CHAPS. For each wash, the resin was incubated at 37°C for 15 minutes prior to allowing the buffer to flow through. Subsequently, the resin was washed with 3 x 10 CVs of wash buffer to remove remaining detergent before elution. Prior to size exclusion chromatography the column and Akta were soaked for 4 h in 0.5M in NaOH before being washed with TBS containing 0.75% CHAPS.

For crystallography, tagless cb3 and HALC3_919 were produced through cleavage at the SNAC tag at the C-terminus of the design. Briefly, cb3- and HALC3_919-SNAC tagged were first bound to Ni-NTA resin and subsequently incubated in 0.1M CHES, 0.1M NaCl, 0.1 M acetone oxime, 0.5 M guanidinium HCl and 2mM NiCl_2_ for 24 hours prior to collecting the tagless proteins from the flow through. Proteins were further purified by size exclusion chromatography as described above.

### Protein biotinylation

MERS-CoV S ectodomain was biotinylated using BirA biotin-protein ligase standard reaction kit (Avidity) as described before*(53)*. Briefly, 20 μM of S ectodomain was incubated overnight at 4°C with 2.5 μg of BirA enzyme in reaction mixtures containing 1X BiomixB, 1X BiomixA and 40 μM BIO200. S ectodomain was further separated from Bir A by SEC using Superose 6 10/300 GL (GE LifeSciences) and concentrated using 100 kDa filters (Amicon).

### Binding assays to biotinylated MERS-CoV RBD *via* yeast-surface display

Yeast display experiments were performed as described previously*(32)*. 25,000 designs were reverse translated and codon optimized for *S. cerevisiae* expression using DNAworks 2.0 and then ordered as 240bp DNA oligos (Agilent) with 5’ GGTGGATCAGGAGGTTCG and 3’ GGAAGCGGTGGAAGTGGG adapters. The SSM libraries were ordered in the same manner. Affinity enhancing mutations identified in the SSM libraries were then ordered as DNA ultramers (IDT) with library diversities of approximately 1E6.

The oligo pools were amplified using Kapa Hifi polymerase (Kapa Biosystems). qPCR reactions were prepared in duplicate 25 μL reactions with the first being run to determine the cycle number for half maximal amplification followed by a second production run. These were then purified from an agarose gel (Zymo Research) and amplified again to obtain sufficient material for transformation. 2 μg of linearized petcon3 and 6ug of insert were transformed into EBY100 through electroporation*(54)*.

Yeast cultures were maintained in C-Trp-Ura media with 2% glucose grown at 30°C. Expression was induced by inoculating 10 ml of SGCAA with 250 μL of C-Trp-Ura culture. Transformed libraries underwent 4 sorts. For all sorts, SGCAA-induced cultures were harvested via centrifugation at 4000xg for 4 minutes. Then the cells were washed once in PBS with 1% BSA (PBSF). The staining occurred for an hour at room temperature followed by 3 PMSF washed in ice cold PBSF. Cells were kept on ice before being resuspended in 50 μL immediately prior to sorting. In sort 1, cells were collected that were FITC positive indicating successful design expression based on anti-myc FITC binding to a myc tag at the C-terminus of the design. In sort two, cells were stained with both anti-myc FITC (Immunology Consultants Laboratory) and streptavidin-PE (SAPE) (Thermo Fisher) preincubated with 1uM biotinylated MERS-CoV S RBD at a 1:4 ratio. Cells that were FITC and PE positive were collected. These cells were grown in C-Trp-Ura for two days and then induced before a second sort following the same protocol was performed. A final titration sort at 1000 nM, 200 nM and 40 nM with RBD and SAPE in a 1:1 ratio was performed to identify the highest affinity binders. Yeast surface display data and determination of SC50s was done using published tools *(32)*.

### Lyophilization

nSB175_cb3 (cb3-GSG-SB175) was buffer exchanged into water and then lyophilized **o**vernight at a concentration of 2 mg/ml using a SP Scientific BenchTop Pro lyophilizer. Lyophilized trimeric miniprotein was stored at room temperature until reconstitution in TBS.

### Measurement of binding affinities using surface plasmon resonance (SPR)

SPR experiments were performed using a Biacore 8K and the biotin CAP chip (Cytiva) for both the monomeric and trimeric designs. For the monomers, approximately 250 RU of biotinylated MERS-CoV S RBD was captured. An 8-step two fold single cycle kinetic dilution series from 500 nM was used to assess affinity with 120 s association and 900 s dissociation at 30 ul/min at 25°C. For trimeric constructs, approximately 250 RU of biotinylated MERS-CoV S protein was captured and an 8-step two-fold single cycle kinetic dilution series from 25 nM was used to assess affinity with 120 s association and 30 min dissociation at 30 ul/min at 25°C. The chip was regenerated between runs with 0.25 M NaOH and 6 M guanidinium hydrochloride. K_D_s were determined through global langmuir 1:1 model fitting using the Biacore Evaluation software.

### Crystallization and structure determination

Freshly purified MERS-CoV S RBD was concentrated to 20 mg/ml, mixed with a 2-fold molar excess of purified miniprotein cb3 and incubated ON at 4 before setting up crystallization screening using a mosquito robot. Crystals were obtained at 22°C by hanging drop vapor diffusion when MERS-CoV-RBD/miniprotein cb3 complex was mixed with mother liquor solution (200 nl final volume) containing: 0.16 M MgCl_2_, 0.08 M Tris-HCl pH 8.5, 24%(w/v) PEG4000, 20% (v/v) glycerol. Crystals were flash frozen in liquid nitrogen using 20 % glycerol as cryoprotectant. Diffraction data were collected at the National Synchrotron Light Source II (Brookhaven) beamline AMX (17-ID-1), and processed with the XDS software package*(55)*. Initial phases were obtained by molecular replacement using Phenix-Phaser*(56)* and the PDB 4L3N*(57)* as a search model. Several subsequent rounds of model building, and refinement were performed using Coot*(58)* and Phenix-Refine*(59)* to arrive at a final model for the binary complex at 1.85 Å resolution in space group P2_1_2_1_2_1_. Model validation was done using Molprobity*(60)* and Privateer*(61)*.

For HALC3_919, the protein was purified from 0.5 L cultures as described above. The sample was then concentrated to 15 mg/ml in 20 mM Tris 50 mM NaCl pH 8. Sitting drop broad screens were set up at 1:1, 1:2, 2:1 ratios of crystallization condition:protein using a mosquito pipetting instrument. JCSG+ (Molecular Dimensions), Morpheus (Molecular Dimensions), PACT premier (Molecular Dimensions), Midas (Molecular Dimensions), Proplex (Molecular Dimensions), Index (Hampton Research). The condition that yielded diffracting crystals was JCSG+ A6 (0.2 M Lithium sulfate monohydrate, 0.1 M Sodium citrate tribasic dihydrate pH 4.2, 20% w/v PEG 1000) cryoprotected with 25% glycerol. Data was collected at the Advanced Photon Source synchrotron, beamline 24ID-C. XDS 20220110*(55)* was used for image integration, scaling and merging with Aimless*(62)*. The structures were phased using molecular replacement with the monomeric design models as search models in Phaser 2.8*(56)*. Models were built with Coot*(58)* and refined with Phenix refine*(63)* and RefMac*(64)* from CCP4 7.1*(62)* suite. Model validation was done using MolProbity 4.5.1*(60)*.

### VSV pseudotyped virus production

MERS-CoV S pseudotyped vesicular stomatitis virus (VSV) corresponding to the different strains were generated as previously described*(53)*. Briefly, HEK293T cells seeded in poly-D-lysine coated 10-cm dishes in DMEM supplemented with 10% FBS and 1% PenStrep were transfected with a mixture of 24 µg of the corresponding plasmids, 60 μL Lipofectamine 2000 (Life Technologies) in 3 ml of Opti-MEM, following manufacturer’s instructions. After 5 h at 37°C, DMEM supplemented with 20% FBS and 2% PenStrep was added. The next day, cells were washed three times with DMEM and were transduced with VSVΔG-luc *(65)*. After 2 h, virus inoculum was removed and cells were washed five times with DMEM prior to the addition of DMEM supplemented with anti-VSV-G antibody [Il-mouse hybridoma supernatant diluted 1 to 25 (v/v), from CRL-2700, ATCC] to minimize parental background. After 18-24 h, supernatants containing pseudotyped VSV were harvested, centrifuged at 2,000 x g for 5 min to remove cellular debris, filtered with a 0,45 µm membrane, concentrated 10 times using a 30 kDa cut off membrane (Amicon), aliquoted, and frozen at −80°C.

### Pseudotyped VSV neutralizations

VeroE6-TMPRSS2 cells were seeded into coated clear bottom white walled 96-well plates at 40,000 cells/well and cultured overnight at 37°C. Eleven 2-fold or 3-fold serial dilutions of the corresponding miniprotein were prepared in DMEM. MERS-CoV S pseudotyped VSV diluted 1 to 20 in DMEM were added 1:1 (v/v) to each miniprotein dilution and mixtures of 50 μL volume were incubated for 45-60 min at 37°C. VeroE6-TMPRSS2 cells were washed three times with DMEM and 40 μL of the mixture containing pseudotyped virus and miniprotein were added. Two hours later, 40 μL of DMEM were added to the cells. After 17-20 h, 70 μL of One-Glo-EX substrate (Promega) were added to each well and incubated on a plate shaker in the dark at 37°C. After 5-15 min incubation, plates were read on a Biotek plate reader. Measurements were done in duplicate or triplicate with at least two biological replicates. Relative luciferase units were plotted and normalized in Prism (GraphPad): cells without pseudotyped virus added were defined as 0 % infection or 100% neutralization, and cells with virus only (no miniprotein) were defined as 100 % infection or 0 % neutralization.

### Western Blot

Pseudotyped VSV particles (15 μL) were mixed with 4X SDS-PAGE loading buffer, run on a 4%–15% gradient Tris-Glycine Gel (BioRad) and transferred to a PVDF membrane using the protocol “mix molecular weight” of the Trans-Blot Turbo System (BioRad). Membranes were blocked with 5 % milk in TBS-T (20 mM Tris-HCl pH 8.0, 150 mM NaCl supplemented with 0.05 % Tween-20) at room temperature and with agitation. After 1 h, the stem-helix specific antibody B6 monoclonal antibody*(48)* was added at 3 µg/ml and incubated overnight at 4°C with agitation. Membranes were washed three times with TBS-T and an Alexa Fluor 680-conjugated goat anti-human secondary antibody (1:50,000 dilution, Jackson ImmunoResearch, 109-625-098) was added and incubated during 1 h at room temperature. Membranes were washed three times with TBS-T after which a LI-COR processor was used to develop the western blot.

### Biolayer interferometry assay (BLI)

Competition binding experiments were performed using BLI on an Octet Red (Sartorius) instrument operated at 30°C with 1000 rpm shaking. Biotinylated MERS-CoV S RBD at 3 µg/mL was immobilized to reach 1 nm shift in undiluted 10X kinetics buffer (Pall) to streptavidin (SA) biosensors (Sartorius) that were pre-hydrated in 10X kinetics buffer for 10 minutes prior the experiment. Biosensors were subsequently dipped into 10x Kinetics buffer to stabilize and remove any unbound protein after which loaded biosensors were dipped into a solution containing either nSB175_cb3 at 0.5 nM in 10X kinetics buffer or only 10X kinetics buffer, until steady state was achieved (1^st^ association step). A 2^nd^ association step was followed by dipping biosensors in a solution containing 0.5 µM nSB175_cb3 and 0.5 µM hDPP4 or only 0.5 µM hDPP4 for 300 seconds. Dissociation phase was followed by dipping the tips into the 10X Kinetics buffer. The data were plotted in Graphpad Prism (v.10.2.2).

### Spike mediated-cell fusion assay

We use a split GFP system to study whether spike mediated-cell fusion is inhibited by mini proteins. Experiments were conducted as previously described in Addetia et al, 2023 with some modifications. In brief, BHK-21-GFP_1-10_ cells were seeded into 6-well plates at a density of 1 x 10^6^ cells per well. The following day, the growth media was removed, cells were washed one time with DMEM, placed in DMEM containing 10% FBS and 1% PenStrep and transfected with 4 µg per well of the corresponding S protein using Lipofectamine 2000. The same day, VeroE6-TMPRSS2-GFP_11_ cells were split into 96-well, glass bottom, black walled plates (CellVis) at a density of 36,000 cells per well. Twenty-four hours after transfection, BHK-21-GFP_1-10_ expressing the S protein were washed three times using FluoroBrite DMEM (Thermo Fisher), detached using an enzyme-free cell dissociation buffer (Gibco), passed through a cell strainer to remove aggregates and incubated at 65,000 cells/ml (for MERS-CoV S Jordan/2012, United Kingdom/2012, Seoul/2015 and Kenya/2019) or at 90,000 cells/ml (for EMC/2012) with different concentrations of miniprotein nSB175_cb3 during 30-45 min at 37°C. VeroE6-TMPRSS2-GFP_11_ cells were washed four times with FluoroBrite DMEM and the transfected BHK-21-GFP_1-10_ cells incubated with the miniprotein were plated on top of it. Cells were incubated at 37°C and 5 % CO_2_ in a Cytation 7 plate Imager (Biotek) and both brightfield and GFP images were collected every 30 minutes for 18 hours. Fusogenicity was assessed by measuring the area showing GFP fluorescence for each image using Gen5 Image Prime v3.11 software.

### Flow cytometry

To measure surface expression of the MERS-CoV S proteins, 1 x 10^6^ transiently transfected BHK-21-GFP_1-10_ cells were collected by centrifugation at 500 x g for 2 min. Cells were washed once with PBS and fixed by incubation with 2% paraformaldehyde during 10-15 min at room temperature. Cells were subsequently washed three times with eBioscience Flow Cytometry Staining Buffer (Invitrogen) and incubated with 4 µg/mL of B6 mAb*(48)*, an S stem-helix directed antibody that recognizes all MERS-CoV S variants, for 1 h in ice. Cells were washed three times with eBioscience Flow Cytometry Staining Buffer and labeled with a PE-conjugated anti-Human IgG Fc antibody (Thermo Fisher) for 1 h in ice. Cells were washed three times and resuspended in 500 μL eBioscience Flow Cytometry Staining Buffer. Labeled cells were analyzed using a BD FACSymphony A3. Cells were gated on singleton events and a total of 10,000 singleton events were collected for each sample. The fraction of S-positive cells was determined in FlowJo 10.8.1 by gating singleton events for the mock transfected cells on PE intensity.

### CryoEM sample preparation, data collection and data processing for MERS-CoV S in complex with nSB175_cb3 (cb3_GSG_SB175) and nSB175_cb3 linker 7

3µl of MERS-CoV S in complex with nSB175_cb3 were loaded at 0.15 mg/ml onto freshly glow-discharged NiTi grids covered with a thin layer of home-made continuous carbon prior to plunge freezing using a vitrobot MarkIV (ThermoFisher Scientific) with a blot force of −1 and 4.5 sec blot time at 100% humidity and 21°C. For MERS-CoV S in complex with nSB175_cb3 linker 7, 3 µl of S at 1.5 mg/ml were loaded onto UltrAufoil*(66)* grids prior to plunge freezing using a vitrobot MarkIV with a blot force of 0 and 6-6.5 sec blot time at 100% humidity and 21°C. For MERS-CoV S in complex with nSB175_cb3 complex and nSB175_cb3 linker 7, 13,530 and 8751 movies were collected, respectively, with a defocus range comprised between −0.8 and −2.0 μm. Both datasets were acquired using a FEI Titan Krios transmission electron microscope operated at 300 kV equipped with a Gatan K3 direct detector and a Gatan Quantum GIF energy filter, operated with a slit width of 20eV. Automated data collection was carried out using the Leginon software*(67)* at a nominal magnification of 105,000x corresponding to a pixel size of 0.843 Å. The dose rate was adjusted to 9 counts/pixel/s, and each movie was acquired in counting mode fractionated in 100 frames of 40 ms. Movie frame alignment, estimation of the microscope contrast-transfer function parameters, particle picking and extraction were carried out using Warp*(68)*. One round of reference-free 2D classification was performed using cryoSPARC*(69)* with binned particles to select well-defined particle images. To further improve particle picking, we trained the Topaz picker*(70)* on the Warp-picked particles belonging to the selected 2D classes. Topaz-picked particles were extracted and 2D classified using cryoSPARC. Topaz-duplicated picked particles were removed using 70 Å (for MERS-CoV S-nSB175_cb3 complex) or 90 Å (for MERS-CoV S-nSB175_cb3 linker 7 complex) as a minimum distance cutoff. Initial model was generated using ab-initio reconstruction in cryoSPARC and used as reference for a non-uniform refinement*(71)* (NUR). Particles were transferred from cryoSPARC to Relion*(72)* using pyem*(73)* (https://github.com/asarnow/pyem) to be subjected to one round of 3D classification with 50 iterations, using the NUR map as a reference model (angular sampling 7.5 ° for 25 iterations and 1.8 ° with local search for 25 iterations) and without imposing symmetry. For MERS-CoV S in complex with nSB175_cb3, we identified two main populations corresponding to trimers with two or with 3 RBDs engaged with the bound miniprotein. Each of the two populations were processed separately after this step. Particles selected were then subjected to NUR with per-particle defocus refinement using cryoSPARC applying C3 symmetry for particles with 3 RBDs engaged by nSB175_cb3 and without symmetry for particles engaging 2 RBDs before carrying out Bayesian polishing*(74)* using Relion. For MERS-CoV S in complex with nSB175_cb3 linker 7, selected particles from Relion 3D classification were subjected to another round of 2D classification followed by a Bayesian polishing*(74)* using Relion. During Bayesian polishing*(74)* particles were re-extracted with a box size of 512 pixels at a pixel size of 1 Å. After polishing, particles from both complexes were subjected to 2D classification in cryoSPARC followed by either NUR with per-particle defocus refinement for MERS-CoV S in complex with nSB175_cb3 or ab-initio and heterogenous refinement to further clean the dataset before NUR with per-particle defocus refinement for MERS-CoV S in complex with nSB175_cb3 linker 7. To improve the density of the MERS-CoV S RBD in complex with nSB175_cb3 linker 7 map, particles were subjected to Relion 3D focused classification without refining angles and shifts using a soft mask encompassing one cb3-bound RBD. Selected particles were subjected in paralell i) to a local refinement using a soft mask encompassing two RBDs with bound cb3s and local resolution estimation using CryoSPARC, which yielded a 3.2 Å resolution map of 2 RBDs in complex with two cb3, and ii) to a Relion 3D focused classification without refining angles and shifts using a soft mask encompassing the less visible RBD and cb3 region (that was not part of the mask used for the first focused 3D classification) followed by NUR with per-particle defocus refinement of the selected particles, that allowed us to obtain a full map in which each RBDs were engaged by one cb3. Reported resolutions are based on the gold-standard Fourier shell correlation using 0.143 criterion*(75)* and Fourier shell correlation curves were corrected for the effects of soft masking by high-resolution noise substitution*(76)*.

### CryoEM model building and refinement

#### *In vivo* experiments

Animal husbandry and all *in vivo* experiments were performed at the University of North Carolina at Chapel Hill with prior approval from the Institutional Animal Care and Use Committee (IACUC protocol: 22-184) in a Biosafety Level 3 (BSL-3) laboratory. Mice were inoculated with either vehicle (TBS), nSB175_cb3 (cb3-GSG-SB175), influenza miniprotein (Flu-HA) monomer*(32)* or were left untreated. The Flu-Ha miniprotein, MSGSQHDEFGKWMIEKLKEAVDRGNKISAEFLYNLGKNFVRNPDILKQMEELRKSLHGSGSHHWGSTHHHHH H, was redesigned to improve its solubility. Twenty-five- to 28-week-old male and female 288/330 mice on a C57BL/6J background were used. Mice were anesthetized with a mixture of xylazine and ketamine prior to nSB175_cb3 treatment (6.25 mg/kg) and viral infection with mouse-adapted MERS-CoV (1×10^5^ PFU) via intranasal instillation. Mice were monitored daily for weiGht loss and siGns of morbidity.At the indicated time post infection, mice were euthanized via isoflurane overdose. Lung tissue was assessed for gross pathology and the inferior lobe was harvested in PBS with glass beads and stored at −80C for viral titer via plaque assay.

## REFERENCES

1. A. M. Zaki, S. van Boheemen, T. M. Bestebroer, A. D. Osterhaus, R. A. Fouchier, Isolation of a novel coronavirus from a man with pneumonia in Saudi Arabia. N. Engl. J. Med. 367, 1814–1820 (2012).

2. A. Bermingham, M. A. Chand, C. S. Brown, E. Aarons, C. Tong, C. Langrish, K. Hoschler, K. Brown, M. Galiano, R. Myers, R. G. Pebody, H. K. Green, N. L. Boddington, R. Gopal, N. Price, W. Newsholme, C. Drosten, R. A. Fouchier, M. Zambon, Severe respiratory illness caused by a novel coronavirus, in a patient transferred to the United Kingdom from the Middle East, September 2012. Euro Surveill. 17, 20290 (2012).

3. MERS-CoV worldwide overviewEuropean Centre for Disease Prevention and Control (2024) (available at https://www.ecdc.europa.eu/en/middle-east-respiratory-syndrome-coronavirus-mers-cov-situation-update).

4. J. S. Sabir, T. T. Lam, M. M. Ahmed, L. Li, Y. Shen, S. E. Abo-Aba, M. I. Qureshi, M. Abu-Zeid, Y. Zhang, M. A. Khiyami, N. S. Alharbi, N. H. Hajrah, M. J. Sabir, M. H. Mutwakil, S. A. Kabli, F. A. Alsulaimany, A. Y. Obaid, B. Zhou, D. K. Smith, E. C. Holmes, H. Zhu, Y. Guan, Co-circulation of three camel coronavirus species and recombination of MERS-CoVs in Saudi Arabia. Science 351, 81–84 (2016).

5. P. M. Munyua, I. Ngere, E. Hunsperger, A. Kochi, P. Amoth, L. Mwasi, S. Tong, A. Mwatondo, N. Thornburg, M.-A. Widdowson, M. K. Njenga, Low-Level Middle East Respiratory Syndrome Coronavirus among Camel Handlers, Kenya, 2019. Emerg. Infect. Dis. 27, 1201–1205 (2021).

6. M. A. Tortorici, D. Veesler, Structural insights into coronavirus entry. Adv. Virus Res. 105, 93–116 (2019).

7. A. C. Walls, X. Xiong, Y. J. Park, M. A. Tortorici, J. Snijder, J. Quispe, E. Cameroni, R. Gopal, M. Dai, A. Lanzavecchia, M. Zambon, F. A. Rey, D. Corti, D. Veesler, Unexpected Receptor Functional Mimicry Elucidates Activation of Coronavirus Fusion. Cell 176, 1026–1039.e15 (2019).

8. A. C. Walls, M. A. Tortorici, B. J. Bosch, B. Frenz, P. J. M. Rottier, F. DiMaio, F. A. Rey, D. Veesler, Cryo-electron microscopy structure of a coronavirus spike glycoprotein trimer. Nature 531, 114–117 (2016).

9. A. Addetia, C. Stewart, A. J. Seo, K. R. Sprouse, A. Y. Asiri, M. Al-Mozaini, Z. A. Memish, A. Alshukairi, D. Veesler, Mapping immunodominant sites on the MERS-CoV spike glycoprotein targeted by infection-elicited antibodies in humansbioRxiv, 2024.03.31.586409 (2024).

10. K. S. Corbett, D. K. Edwards, S. R. Leist, O. M. Abiona, S. Boyoglu-Barnum, R. A. Gillespie, S. Himansu, A. Schäfer, C. T. Ziwawo, A. T. DiPiazza, K. H. Dinnon, S. M. Elbashir, C. A. Shaw, A. Woods, E. J. Fritch, D. R. Martinez, K. W. Bock, M. Minai, B. M. Nagata, G. B. Hutchinson, K. Wu, C. Henry, K. Bahl, D. Garcia-Dominguez, L. Ma, I. Renzi, W.-P. Kong, S. D. Schmidt, L. Wang, Y. Zhang, E. Phung, L. A. Chang, R. J. Loomis, N. E. Altaras, E. Narayanan, M. Metkar, V. Presnyak, C. Liu, M. K. Louder, W. Shi, K. Leung, E. S. Yang, A. West, K. L. Gully, L. J. Stevens, N. Wang, D. Wrapp, N. A. Doria-Rose, G. Stewart-Jones, H. Bennett, G. S. Alvarado, M. C. Nason, T. J. Ruckwardt, J. S. McLellan, M. R. Denison, J. D. Chappell, I. N. Moore, K. M. Morabito, J. R. Mascola, R. S. Baric, A. Carfi, B. S. Graham, SARS-CoV-2 mRNA vaccine design enabled by prototype pathogen preparedness. Nature 586, 567–571 (2020).

11. D. Corti, L. A. Purcell, G. Snell, D. Veesler, Tackling COVID-19 with neutralizing monoclonal antibodies. Cell (2021), doi:10.1016/j.cell.2021.05.005.

12. V. S. Raj, H. Mou, S. L. Smits, D. H. W. Dekkers, M. A. Müller, R. Dijkman, D. Muth, J. A. A. Demmers, A. Zaki, R. A. M. Fouchier, V. Thiel, C. Drosten, P. J. M. Rottier, A. D. M. E. Osterhaus, B. J. Bosch, B. L. Haagmans, Dipeptidyl peptidase 4 is a functional receptor for the emerging human coronavirus-EMC. Nature 495, 251–254 (2013).

13. Y.-J. Park, A. C. Walls, Z. Wang, M. M. Sauer, W. Li, M. A. Tortorici, B.-J. Bosch, F. DiMaio, D. Veesler, Structures of MERS-CoV spike glycoprotein in complex with sialoside attachment receptors. Nat. Struct. Mol. Biol. 26, 1151–1157 (2019).

14. W. Li, R. J. G. Hulswit, I. Widjaja, V. S. Raj, R. McBride, W. Peng, W. Widagdo, M. A. Tortorici, B. van Dieren, Y. Lang, J. W. M. van Lent, J. C. Paulson, C. A. M. de Haan, R. J. de Groot, F. J. M. van Kuppeveld, B. L. Haagmans, B.-J. Bosch, Identification of sialic acid-binding function for the Middle East respiratory syndrome coronavirus spike glycoprotein. Proceedings of the National Academy of Sciences 114, E8508–E8517 (2017).

15. G. Lu, Y. Hu, Q. Wang, J. Qi, F. Gao, Y. Li, Y. Zhang, W. Zhang, Y. Yuan, J. Bao, B. Zhang, Y. Shi, J. Yan, G. F. Gao, Molecular basis of binding between novel human coronavirus MERS-CoV and its receptor CD26. Nature 500, 227–231 (2013).

16. J. Pallesen, N. Wang, K. S. Corbett, D. Wrapp, R. N. Kirchdoerfer, H. L. Turner, C. A. Cottrell, M. M. Becker, L. Wang, W. Shi, W.-P. Kong, E. L. Andres, A. N. Kettenbach, M. R. Denison, J. D. Chappell, B. S. Graham, A. B. Ward, J. S. McLellan, Immunogenicity and structures of a rationally designed prefusion MERS-CoV spike antigen. Proc. Natl. Acad. Sci. U. S. A. 114, E7348–E7357 (2017).

17. Y. Yuan, D. Cao, Y. Zhang, J. Ma, J. Qi, Q. Wang, G. Lu, Y. Wu, J. Yan, Y. Shi, X. Zhang, G. F. Gao, Cryo-EM structures of MERS-CoV and SARS-CoV spike glycoproteins reveal the dynamic receptor binding domains. Nat. Commun. 8, 15092 (2017).

18. W. Widagdo, V. S. Raj, D. Schipper, K. Kolijn, G. van Leenders, B. J. Bosch, A. Bensaid, J. Segales, W. Baumgartner, A. Osterhaus, M. P. Koopmans, J. M. A. van den Brand, B. L. Haagmans, Differential Expression of the Middle East Respiratory Syndrome Coronavirus Receptor in the Upper Respiratory Tracts of Humans and Dromedary Camels. J. Virol. 90, 4838–4842 (2016).

19. Q. Xiong, L. Cao, C. Ma, M. A. Tortorici, C. Liu, J. Si, P. Liu, M. Gu, A. C. Walls, C. Wang, L. Shi, F. Tong, M. Huang, J. Li, C. Zhao, C. Shen, Y. Chen, H. Zhao, K. Lan, D. Corti, D. Veesler, X. Wang, H. Yan, Close relatives of MERS-CoV in bats use ACE2 as their functional receptors. Nature 612, 748–757 (2022).

20. V. D. Menachery, K. H. Dinnon 3rd, B. L. Yount Jr, E. T. McAnarney, L. E. Gralinski, A. Hale, R. L. Graham, T. Scobey, S. J. Anthony, L. Wang, B. Graham, S. H. Randell, W. I. Lipkin, R. S. Baric, Trypsin treatment unlocks barrier for zoonotic bat Coronavirus infection. J. Virol. 94 (2020), doi:10.1128/JVI.01774-19.

21. L. V. Tse, Y. J. Hou, E. McFadden, R. E. Lee, T. D. Scobey, S. R. Leist, D. R. Martinez, R. M. Meganck, A. Schäfer, B. L. Yount, T. Mascenik, J. M. Powers, S. H. Randell, Y. Zhang, L. Wang, J. Mascola, J. S. McLellan, R. S. Baric, A MERS-CoV antibody neutralizes a pre-emerging group 2c bat coronavirus. Sci. Transl. Med. 15, eadg5567 (2023).

22. C.-B. Ma, C. Liu, Y.-J. Park, J. Tang, J. Chen, Q. Xiong, J. Lee, C. Stewart, D. Asarnow, J. Brown, M. Alejandra Tortorici, X. Yang, Y.-H. Sun, Y.-M. Chen, X. Yu, J.-Y. Si, P. Liu, F. Tong, M.-L. Huang, J. Li, Z.-L. Shi, Z. Deng, D. Veesler, H. Yan, Multiple independent acquisitions of ACE2 usage in MERS-related coronavirusesbioRxiv, 2023.10.02.560486 (2024).

23. Y.-J. Park, C. Liu, J. Lee, J. T. Brown, C.-B. Ma, P. Liu, Q. Xiong, C. Stewart, A. Addetia, C. J. Craig, M. Alejandra Tortorici, A. Alshukari, T. Starr, H. Yan, D. Veesler, Molecular basis of convergent evolution of ACE2 receptor utilization among HKU5 coronavirusesbioRxiv, 2024.08.28.608351 (2024).

24. A. C. Hunt, J. B. Case, Y.-J. Park, L. Cao, K. Wu, A. C. Walls, Z. Liu, J. E. Bowen, H.-W. Yeh, S. Saini, L. Helms, Y. T. Zhao, T.-Y. Hsiang, T. N. Starr, I. Goreshnik, L. Kozodoy, L. Carter, R. Ravichandran, L. B. Green, W. L. Matochko, C. A. Thomson, B. Vögeli, A. Krüger, L. A. VanBlargan, R. E. Chen, B. Ying, A. L. Bailey, N. M. Kafai, S. E. Boyken, A. Ljubetič, N. Edman, G. Ueda, C. M. Chow, M. Johnson, A. Addetia, M. J. Navarro, N. Panpradist, M. Gale, B. S. Freedman, J. D. Bloom, H. Ruohola-Baker, S. P. J. Whelan, L. Stewart, M. S. Diamond, D. Veesler, M. C. Jewett, D. Baker, Multivalent designed proteins neutralize SARS-CoV-2 variants of concern and confer protection against infection in mice. Sci. Transl. Med. 0, eabn1252.

25. L. Cao, I. Goreshnik, B. Coventry, J. B. Case, L. Miller, L. Kozodoy, R. E. Chen, L. Carter, A. C. Walls, Y. J. Park, E. M. Strauch, L. Stewart, M. S. Diamond, D. Veesler, D. Baker, De novo design of picomolar SARS-CoV-2 miniprotein inhibitors. Science 370, 426–431 (2020).

26. J. B. Case, R. E. Chen, L. Cao, B. Ying, E. S. Winkler, M. Johnson, I. Goreshnik, M. N. Pham, S. Shrihari, N. M. Kafai, A. L. Bailey, X. Xie, P.-Y. Shi, R. Ravichandran, L. Carter, L. Stewart, D. Baker, M. S. Diamond, Ultrapotent miniproteins targeting the SARS-CoV-2 receptor-binding domain protect against infection and disease. Cell Host Microbe 29, 1151–1161.e5 (2021).

27. J. Lee, J. B. Case, Y.-J. Park, R. Ravichandran, D. Asarnow, M. A. Tortorici, J. T. Brown, S. Sanapala, L. Carter, D. Baker, M. S. Diamond, D. Veesler, A pan-variant miniprotein inhibitor protects against SARS-CoV-2 variants. bioRxiv (2024), doi:10.1101/2024.08.08.606885.

28. J. E. Bowen, Y.-J. Park, C. Stewart, J. T. Brown, W. K. Sharkey, A. C. Walls, A. Joshi, K. R. Sprouse, M. McCallum, M. A. Tortorici, N. M. Franko, J. K. Logue, I. G. Mazzitelli, A. W. Nguyen, R. P. Silva, Y. Huang, J. S. Low, J. Jerak, S. W. Tiles, K. Ahmed, A. Shariq, J. M. Dan, Z. Zhang, D. Weiskopf, A. Sette, G. Snell, C. M. Posavad, N. T. Iqbal, J. Geffner, A. Bandera, A. Gori, F. Sallusto, J. A. Maynard, S. Crotty, W. C. Van Voorhis, C. Simmerling, R. Grifantini, H. Y. Chu, D. Corti, D. Veesler, SARS-CoV-2 spike conformation determines plasma neutralizing activity elicited by a wide panel of human vaccines. Sci Immunol 7, eadf1421 (2022).

29. L. Piccoli, Y. J. Park, M. A. Tortorici, N. Czudnochowski, A. C. Walls, M. Beltramello, C. Silacci-Fregni, D. Pinto, L. E. Rosen, J. E. Bowen, O. J. Acton, S. Jaconi, B. Guarino, A. Minola, F. Zatta, N. Sprugasci, J. Bassi, A. Peter, A. De Marco, J. C. Nix, F. Mele, S. Jovic, B. F. Rodriguez, S. V. Gupta, F. Jin, G. Piumatti, G. Lo Presti, A. F. Pellanda, M. Biggiogero, M. Tarkowski, M. S. Pizzuto, E. Cameroni, C. Havenar-Daughton, M. Smithey, D. Hong, V. Lepori, E. Albanese, A. Ceschi, E. Bernasconi, L. Elzi, P. Ferrari, C. Garzoni, A. Riva, G. Snell, F. Sallusto, K. Fink, H. W. Virgin, A. Lanzavecchia, D. Corti, D. Veesler, Mapping Neutralizing and Immunodominant Sites on the SARS-CoV-2 Spike Receptor-Binding Domain by Structure-Guided High-Resolution Serology. Cell 183, 1024–1042.e21 (2020).

30. A. J. Greaney, A. N. Loes, L. E. Gentles, K. H. D. Crawford, T. N. Starr, K. D. Malone, H. Y. Chu, J. D. Bloom, Antibodies elicited by mRNA-1273 vaccination bind more broadly to the receptor binding domain than do those from SARS-CoV-2 infection. Sci. Transl. Med. 13 (2021), doi:10.1126/scitranslmed.abi9915.

31. J. Dou, A. A. Vorobieva, W. Sheffler, L. A. Doyle, H. Park, M. J. Bick, B. Mao, G. W. Foight, M. Y. Lee, L. A. Gagnon, L. Carter, B. Sankaran, S. Ovchinnikov, E. Marcos, P.-S. Huang, J. C. Vaughan, B. L. Stoddard, D. Baker, De novo design of a fluorescence-activating β-barrel. Nature 561, 485–491 (2018).

32. L. Cao, B. Coventry, I. Goreshnik, B. Huang, W. Sheffler, J. S. Park, K. M. Jude, I. Marković, R. U. Kadam, K. H. G. Verschueren, K. Verstraete, S. T. R. Walsh, N. Bennett, A. Phal, A. Yang, L. Kozodoy, M. DeWitt, L. Picton, L. Miller, E.-M. Strauch, N. D. DeBouver, A. Pires, A. K. Bera, S. Halabiya, B. Hammerson, W. Yang, S. Bernard, L. Stewart, I. A. Wilson, H. Ruohola-Baker, J. Schlessinger, S. Lee, S. N. Savvides, K. C. Garcia, D. Baker, Design of protein-binding proteins from the target structure alone. Nature 605, 551–560 (2022).

33. I. Ngere, E. A. Hunsperger, S. Tong, J. Oyugi, W. Jaoko, J. L. Harcourt, N. J. Thornburg, H. Oyas, M. Muturi, E. M. Osoro, J. Gachohi, C. Ombok, J. Dawa, Y. Tao, J. Zhang, L. Mwasi, C. Ochieng, A. Mwatondo, B. Bodha, D. Langat, A. Herman-Roloff, M. K. Njenga, M.-A. Widdowson, P. M. Munyua, Outbreak of Middle East Respiratory Syndrome Coronavirus in Camels and Probable Spillover Infection to Humans in Kenya. Viruses 14 (2022), doi:10.3390/v14081743.

34. W. Tai, Y. Wang, C. A. Fett, G. Zhao, F. Li, S. Perlman, S. Jiang, Y. Zhou, L. Du, Recombinant Receptor-Binding Domains of Multiple Middle East Respiratory Syndrome Coronaviruses (MERS-CoVs) Induce Cross-Neutralizing Antibodies against Divergent Human and Camel MERS-CoVs and Antibody Escape Mutants. J. Virol. 91 (2017), doi:10.1128/JVI.01651-16.

35. T.-H. Jang, W.-J. Park, H. Lee, H.-M. Woo, S.-Y. Lee, K.-C. Kim, S. S. Kim, E. Hong, J. Song, J.-Y. Lee, The structure of a novel antibody against the spike protein inhibits Middle East respiratory syndrome coronavirus infections. Sci. Rep. 12, 1260 (2022).

36. H. Kleine-Weber, M. T. Elzayat, L. Wang, B. S. Graham, M. A. Müller, C. Drosten, S. Pöhlmann, M. Hoffmann, Mutations in the Spike Protein of Middle East Respiratory Syndrome Coronavirus Transmitted in Korea Increase Resistance to Antibody-Mediated Neutralization. J. Virol. 93 (2019), doi:10.1128/JVI.01381-18.

37. B. I. M. Wicky, L. F. Milles, A. Courbet, R. J. Ragotte, J. Dauparas, E. Kinfu, S. Tipps, R. D. Kibler, M. Baek, F. DiMaio, X. Li, L. Carter, A. Kang, H. Nguyen, A. K. Bera, D. Baker, Hallucinating symmetric protein assemblies. Science 378, 56–61 (2022).

38. D. A. Vishali, J. Monisha, S. K. Sivakamasundari, J. A. Moses, C. Anandharamakrishnan, Spray freeze drying: Emerging applications in drug delivery. J. Control. Release 300, 93–101 (2019).

39. W. Wang, Lyophilization and development of solid protein pharmaceuticals. Int. J. Pharm. 203, 1–60 (2000).

40. J. E. Bowen, A. Addetia, H. V. Dang, C. Stewart, J. T. Brown, W. K. Sharkey, K. R. Sprouse, A. C. Walls, I. G. Mazzitelli, J. K. Logue, N. M. Franko, N. Czudnochowski, A. E. Powell, E. Dellota Jr, K. Ahmed, A. S. Ansari, E. Cameroni, A. Gori, A. Bandera, C. M. Posavad, J. M. Dan, Z. Zhang, D. Weiskopf, A. Sette, S. Crotty, N. T. Iqbal, D. Corti, J. Geffner, G. Snell, R. Grifantini, H. Y. Chu, D. Veesler, Omicron spike function and neutralizing activity elicited by a comprehensive panel of vaccines. Science 377, 890–894 (2022).

41. A. S. Cockrell, B. L. Yount, T. Scobey, K. Jensen, M. Douglas, A. Beall, X.-C. Tang, W. A. Marasco, M. T. Heise, R. S. Baric, A mouse model for MERS coronavirus-induced acute respiratory distress syndrome. Nat. Microbiol. 2, 16226 (2016).

42. J. C. Almagro, G. Mellado-Sánchez, M. Pedraza-Escalona, S. M. Pérez-Tapia, Evolution of Anti-SARS-CoV-2 Therapeutic Antibodies. Int. J. Mol. Sci. 23 (2022), doi:10.3390/ijms23179763.

43. E. Cameroni, J. E. Bowen, L. E. Rosen, C. Saliba, S. K. Zepeda, K. Culap, D. Pinto, L. A. VanBlargan, A. De Marco, J. di Iulio, F. Zatta, H. Kaiser, J. Noack, N. Farhat, N. Czudnochowski, C. Havenar-Daughton, K. R. Sprouse, J. R. Dillen, A. E. Powell, A. Chen, C. Maher, L. Yin, D. Sun, L. Soriaga, J. Bassi, C. Silacci-Fregni, C. Gustafsson, N. M. Franko, J. Logue, N. T. Iqbal, I. Mazzitelli, J. Geffner, R. Grifantini, H. Chu, A. Gori, A. Riva, O. Giannini, A. Ceschi, P. Ferrari, P. E. Cippà, A. Franzetti-Pellanda, C. Garzoni, P. J. Halfmann, Y. Kawaoka, C. Hebner, L. A. Purcell, L. Piccoli, M. S. Pizzuto, A. C. Walls, M. S. Diamond, A. Telenti, H. W. Virgin, A. Lanzavecchia, G. Snell, D. Veesler, D. Corti, Broadly neutralizing antibodies overcome SARS-CoV-2 Omicron antigenic shift. Nature 602, 664–670 (2022).

44. M. McCallum, N. Czudnochowski, L. E. Rosen, S. K. Zepeda, J. E. Bowen, A. C. Walls, K. Hauser, A. Joshi, C. Stewart, J. R. Dillen, A. E. Powell, T. I. Croll, J. Nix, H. W. Virgin, D. Corti, G. Snell, D. Veesler, Structural basis of SARS-CoV-2 Omicron immune evasion and receptor engagement. Science, eabn8652 (2022).

45. A. C. Walls, K. R. Sprouse, J. E. Bowen, A. Joshi, N. Franko, M. J. Navarro, C. Stewart, E. Cameroni, M. McCallum, E. A. Goecker, E. J. Degli-Angeli, J. Logue, A. Greninger, D. Corti, H. Y. Chu, D. Veesler, SARS-CoV-2 breakthrough infections elicit potent, broad, and durable neutralizing antibody responses. Cell (2022), doi:10.1016/j.cell.2022.01.011.

46. M. McCallum, A. C. Walls, K. R. Sprouse, J. E. Bowen, L. E. Rosen, H. V. Dang, A. De Marco, N. Franko, S. W. Tilles, J. Logue, M. C. Miranda, M. Ahlrichs, L. Carter, G. Snell, M. S. Pizzuto, H. Y. Chu, W. C. Van Voorhis, D. Corti, D. Veesler, Molecular basis of immune evasion by the Delta and Kappa SARS-CoV-2 variants. Science, eabl8506 (2021).

47. M. McCallum, J. Bassi, A. De Marco, A. Chen, A. C. Walls, J. Di Iulio, M. A. Tortorici, M.-J. Navarro, C. Silacci-Fregni, C. Saliba, K. R. Sprouse, M. Agostini, D. Pinto, K. Culap, S. Bianchi, S. Jaconi, E. Cameroni, J. E. Bowen, S. W. Tilles, M. S. Pizzuto, S. B. Guastalla, G. Bona, A. F. Pellanda, C. Garzoni, W. C. Van Voorhis, L. E. Rosen, G. Snell, A. Telenti, H. W. Virgin, L. Piccoli, D. Corti, D. Veesler, SARS-CoV-2 immune evasion by the B.1.427/B.1.429 variant of concern. Science (2021), doi:10.1126/science.abi7994.

48. M. M. Sauer, M. A. Tortorici, Y.-J. Park, A. C. Walls, L. Homad, O. J. Acton, J. E. Bowen, C. Wang, X. Xiong, W. de van der Schueren, J. Quispe, B. G. Hoffstrom, B.-J. Bosch, A. T. McGuire, D. Veesler, Structural basis for broad coronavirus neutralization. Nat. Struct. Mol. Biol. 28, 478–486 (2021).

49. B. Dang, M. Mravic, H. Hu, N. Schmidt, B. Mensa, W. F. DeGrado, SNAC-tag for sequence-specific chemical protein cleavage. Nat. Methods 16, 319–322 (2019).

50. D. Schneidman-Duhovny, Y. Inbar, R. Nussinov, H. J. Wolfson, PatchDock and SymmDock: servers for rigid and symmetric docking. Nucleic Acids Res. 33, W363–7 (2005).

51. T. J. Brunette, M. J. Bick, J. M. Hansen, C. M. Chow, J. M. Kollman, D. Baker, Modular repeat protein sculpting using rigid helical junctions. Proc. Natl. Acad. Sci. U. S. A. 117, 8870–8875 (2020).

52. C. Norn, B. I. M. Wicky, D. Juergens, S. Liu, D. Kim, D. Tischer, B. Koepnick, I. Anishchenko, Foldit Players, D. Baker, S. Ovchinnikov, Protein sequence design by conformational landscape optimization. Proc. Natl. Acad. Sci. U. S. A. 118 (2021), doi:10.1073/pnas.2017228118.

53. M. A. Tortorici, A. C. Walls, A. Joshi, Y.-J. Park, R. T. Eguia, M. C. Miranda, E. Kepl, A. Dosey, T. Stevens-Ayers, M. J. Boeckh, A. Telenti, A. Lanzavecchia, N. P. King, D. Corti, J. D. Bloom, D. Veesler, Structure, receptor recognition, and antigenicity of the human coronavirus CCoV-HuPn-2018 spike glycoprotein. Cell 185, 2279–2291.e17 (2022).

54. L. Benatuil, J. M. Perez, J. Belk, C.-M. Hsieh, An improved yeast transformation method for the generation of very large human antibody libraries. Protein Eng. Des. Sel. 23, 155–159 (2010).

55. W. Kabsch, XDS. Acta Crystallogr. D Biol. Crystallogr. 66, 125–132 (2010).

56. A. J. McCoy, R. W. Grosse-Kunstleve, P. D. Adams, M. D. Winn, L. C. Storoni, R. J. Read, Phaser crystallographic software. J. Appl. Crystallogr. 40, 658–674 (2007).

57. Y. Chen, K. R. Rajashankar, Y. Yang, S. S. Agnihothram, C. Liu, Y.-L. Lin, R. S. Baric, F. Li, Crystal structure of the receptor-binding domain from newly emerged Middle East respiratory syndrome coronavirus. J. Virol. 87, 10777–10783 (2013).

58. P. Emsley, B. Lohkamp, W. G. Scott, K. Cowtan, Features and development of Coot. Acta Crystallogr. D Biol. Crystallogr. 66, 486–501 (2010).

59. D. Liebschner, P. V. Afonine, M. L. Baker, G. Bunkóczi, V. B. Chen, T. I. Croll, B. Hintze, L. W. Hung, S. Jain, A. J. McCoy, N. W. Moriarty, R. D. Oeffner, B. K. Poon, M. G. Prisant, R. J. Read, J. S. Richardson, D. C. Richardson, M. D. Sammito, O. V. Sobolev, D. H. Stockwell, T. C. Terwilliger, A. G. Urzhumtsev, L. L. Videau, C. J. Williams, P. D. Adams, Macromolecular structure determination using X-rays, neutrons and electrons: recent developments in Phenix. Acta Crystallogr D Struct Biol 75, 861–877 (2019).

60. C. J. Williams, J. J. Headd, N. W. Moriarty, M. G. Prisant, L. L. Videau, L. N. Deis, V. Verma, D. A. Keedy, B. J. Hintze, V. B. Chen, S. Jain, S. M. Lewis, W. B. Arendall 3rd, J. Snoeyink, P. D. Adams, S. C. Lovell, J. S. Richardson, D. C. Richardson, MolProbity: More and better reference data for improved all-atom structure validation. Protein Sci. 27, 293–315 (2018).

61. J. Agirre, J. Iglesias-Fernández, C. Rovira, G. J. Davies, K. S. Wilson, K. D. Cowtan, Privateer: software for the conformational validation of carbohydrate structures. Nat. Struct. Mol. Biol. 22, 833–834 (2015).

62. M. D. Winn, C. C. Ballard, K. D. Cowtan, E. J. Dodson, P. Emsley, P. R. Evans, R. M. Keegan, E. B. Krissinel, A. G. W. Leslie, A. McCoy, S. J. McNicholas, G. N. Murshudov, N. S. Pannu, E. A. Potterton, H. R. Powell, R. J. Read, A. Vagin, K. S. Wilson, Overview of the CCP4 suite and current developments. Acta Crystallogr. D Biol. Crystallogr. 67, 235–242 (2011).

63. P. D. Adams, P. V. Afonine, G. Bunkóczi, V. B. Chen, I. W. Davis, N. Echols, J. J. Headd, L.-W. Hung, G. J. Kapral, R. W. Grosse-Kunstleve, A. J. McCoy, N. W. Moriarty, R. Oeffner, R. J. Read, D. C. Richardson, J. S. Richardson, T. C. Terwilliger, P. H. Zwart, PHENIX: a comprehensive Python-based system for macromolecular structure solution. Acta Crystallogr. D Biol. Crystallogr. 66, 213–221 (2010).

64. G. N. Murshudov, A. A. Vagin, E. J. Dodson, Refinement of macromolecular structures by the maximum-likelihood method. Acta Crystallogr. D Biol. Crystallogr. 53, 240–255 (1997).

65. Y. Kaname, H. Tani, C. Kataoka, M. Shiokawa, S. Taguwa, T. Abe, K. Moriishi, T. Kinoshita, Y. Matsuura, Acquisition of complement resistance through incorporation of CD55/decay-accelerating factor into viral particles bearing baculovirus GP64. J. Virol. 84, 3210–3219 (2010).

66. C. J. Russo, L. A. Passmore, Electron microscopy: Ultrastable gold substrates for electron cryomicroscopy. Science 346, 1377–1380 (2014).

67. C. Suloway, J. Pulokas, D. Fellmann, A. Cheng, F. Guerra, J. Quispe, S. Stagg, C. S. Potter, B. Carragher, Automated molecular microscopy: the new Leginon system. J. Struct. Biol. 151, 41–60 (2005).

68. D. Tegunov, P. Cramer, Real-time cryo-electron microscopy data preprocessing with Warp. Nat. Methods 16, 1146–1152 (2019).

69. A. Punjani, J. L. Rubinstein, D. J. Fleet, M. A. Brubaker, cryoSPARC: algorithms for rapid unsupervised cryo-EM structure determination. Nat. Methods 14, 290–296 (2017).

70. T. Bepler, K. Kelley, A. J. Noble, B. Berger, Topaz-Denoise: general deep denoising models for cryoEM and cryoET. Nat. Commun. 11, 5208 (2020).

71. A. Punjani, H. Zhang, D. J. Fleet, Non-uniform refinement: adaptive regularization improves single-particle cryo-EM reconstruction. Nat. Methods 17, 1214–1221 (2020).

72. J. Zivanov, T. Nakane, B. O. Forsberg, D. Kimanius, W. J. Hagen, E. Lindahl, S. H. Scheres, New tools for automated high-resolution cryo-EM structure determination in RELION-3. Elife 7 (2018), doi:10.7554/eLife.42166.

73. D. Asarnow, E. Palovcak, Y. Cheng, UCSF pyem v0. 5. Zenodo 10.5281/zenodo 3576630, 2019 (2019).

74. J. Zivanov, T. Nakane, S. H. W. Scheres, A Bayesian approach to beam-induced motion correction in cryo-EM single-particle analysis. IUCrJ 6, 5–17 (2019).

75. P. B. Rosenthal, R. Henderson, Optimal determination of particle orientation, absolute hand, and contrast loss in single-particle electron cryomicroscopy. J. Mol. Biol. 333, 721–745 (2003).

76. S. Chen, G. McMullan, A. R. Faruqi, G. N. Murshudov, J. M. Short, S. H. Scheres, R. Henderson, High-resolution noise substitution to measure overfitting and validate resolution in 3D structure determination by single particle electron cryomicroscopy. Ultramicroscopy 135, 24–35 (2013).

